# Modeling and validation of parallel co-flow layer widths of viscous liquids in open-capillary trigger valve systems

**DOI:** 10.64898/2026.06.25.734354

**Authors:** Jodie C. Tokihiro, TJ Caira, Ariel Lin, Alya Shaposhnikov, Jamison M. Whitten, Celeste S. Castaneda-Lopez, Xiaojing Su, Albert Shin, Ingrid H. Robertson, Tristan M. Nicholson, Ayokunle O. Olanrewaju, Erwin Berthier, Jean Berthier, Ashleigh B. Theberge

## Abstract

Control of fluids is a hallmark of microfluidic systems and fundamental for the successful application of microfluidic devices. Trigger valves use geometric features to autonomously control the release of fluids in microfluidic devices. Our previous work has adapted geometries used in closed trigger valve systems to enable use in open systems, allowing for open microfluidic devices with up to three trigger valves. Here, we focus on the parallel co-flows produced by sequential release of trigger valves and present a model that predicts their layer widths as a function of the geometric characteristics of the different side channels of each trigger valve. We show layered co-flows with widths as low as 50 microns. Additionally, we expand the use of trigger valves in open microfluidic devices by incorporating 1) varied step heights, 2) devices with up to seven trigger valves, and 3) use of varied fluids and plastics including an application with fibroblasts in collagen. To validate the implementation and use of these trigger valves in open systems, we have developed a theoretical framework to compare predicted outcomes (i.e., fluid travel distance, velocity, and layering width) with our experimental values. This theoretical work offers applications in various fields, including hydrogel patterning for 3D cell culture, organ-on-a-chip models, at-home sample preparation, and autonomous microfluidic systems for biosensing.

## Introduction

Microfluidics is a popular avenue for small-scale experiments such as chemical reactions and cell culture through its ability to move fluids through channels with at least one dimension that is one millimeter or smaller. One of the many branches of microfluidics is capillary microfluidics which has been widely adopted across various disciplines in recent years and uses surface tension to drive microfluidic flow.^1–3^ The field of open capillary-driven microfluidics goes further wherein devices are created through the removal of one or more channel walls conferring additional benefits to their closed-channel counterparts.^4–8^ Open channels offer a multitude of benefits including easy fabrication, simple operation (only requiring the fluid of choice to be added in the inlet reservoir to begin flow), direct access to channel contents, and a wide variety of usable fabrication materials.^9,5,6,10,11^

Within open capillary microfluidic devices, trigger valves–abbreviated henceforth as TGV– are one method for on-chip fluid control and provide a passive and simple approach to fluid control in microfluidic devices, enabling the preprogramming and activation of precisely timed sequences of events on-chip, without any pumps or actuators.^12–15^ TGVs by design are modified passive stop valves that rely on the loss of capillary pressure to pin a fluid at a perpendicular intersection. To release the flow, a miscible fluid flows orthogonal to the TGV, creating two streams of parallel coflows in the main channel. Previously, we presented a mathematical model for trigger valves in open-channel configurations with a comparison to in-lab flow experiments.^16^ Our study was the first step towards understanding the fluid dynamics underlying the microfluidic flow by calculating travel distance of the front meniscus in the main channel as a function of time as each trigger valve is released. One of the benefits of trigger valve systems is the simplicity in operation which relies on fluid pinning (stoppage) at an interface. When desired, an orthogonal flow of miscible liquid is flowed, “triggering” the release of the stopped fluid.

Trigger valves have been used for a variety of applications such as wearable biosensors^17^, electrochemical assays^18^, and one-step immunoassays^19^. There are many different types of geometrical trigger valves that only rely on channel architecture for fluid control including (1) one-level trigger valves, where the trigger valve and destination channel are at the same depth and (2) stair-step trigger valves, where the trigger valve is shallower than the destination channel. Stair-step trigger valves have shown to be a stable approach and is the technique used in this _work._20,21

Adding onto the utilities of TGV systems, is the ability to create layered co-flows–or thin layers of liquids flowing in parallel–within microfluidic channels that are distinct and stable (i.e., no mixing) for a period of time after release in capillary channels. Traditionally, these types of flows are seen in closed-channel microfluidic devices, which offer a wide variety of methods for generating parallel laminar flows. Forced flows are one of the methods for creating streams of fluids conveying benefits to precision in fluid control, time scale, and sub-millimeter size scales.^22–25^ These types of layered flows can also be produced through capillary channel architectures. Jang *et al.* fabricated Y-shaped capillary channels in double sided adhesive and transparency films to create two streams of coflowing fluids–also called stratified streams– with fluid controlled through geometry modifications.^22^ The Walsh group also used a Y-shaped channel structure to generate chemical gradient layers through fluid-filled walls of capillary channels.^25^ More complex networks of side-by-side laminar flows have also been developed for the generation of gradients in microfluidic chips.^26^ Another geometrical feature used to generate parallel layers is bifurcations in which successive bifurcations are used to create a gradient of cells and material cues in hydrogels.^27^ Further, stratified streams of fluids in microfluidic chips have been used for applications such as electrochemical cells^28^, chemical gradient modeling^29^, biofilm development^30^, drug exposure profiling^31^, cell migration^32^, cell patterning^33^, and chemotaxis studies^34^, reactive mixing^35^, and polymer films^23,36–39^.

We present an extension of our previous work on open capillary trigger valves^16^ where the focus was predicting timing of the meniscus fluid front, by now shifting our focus to predicting the width of the parallel layers produced by sequential TGV release. As previously mentioned, these parallel layers have many different applications but a key advantage to this method is the open nature of the device. Due to this there is greater accessibility, multiple fabrication methods and simpler sample control. Here we introduce a model using the geometrical characteristics of the side channels for predicting layer widths of each liquid released by the TGV. In this study, parallel layers down to 50 microns have been achieved, and we demonstrate the successive release of up to seven trigger valves with a singular device. We calculate the pressure at each TGV node and provide a deeper investigation into the contributing flow rates of TGVs resulting in parallel co-flows. This work also compares theoretical layer widths with experimentally-obtained widths using dyed nonanol/aqueous solutions as our flowing liquid and poly(methyl)methacrylate (PMMA)/polystyrene (PS) devices containing five or more trigger valves or differing step heights. This foundational work in predicting fluid layering opens potential applications in a variety of areas such as hydrogel patterning for *in vitro* tissue engineering and thin polymer film generation. To this effect, we also present a proof-of-concept hydrogel patterning application of our TGV system with 3T3 fibroblasts and a collagen-based extracellular matrix.

## Experimental Section

### Device design and fabrication

In this study, five TGV devices were designed, containing a varying amount of TGVs and different cross sections. Table 1 presents critical characteristics of each device. These devices all included a main channel with a cross-section (width by height) of 1000 by 1000 µm. The side channels were oriented along the same side of the device. All side channels had cross-sections with a width of 400 µm and varying heights (step heights) with a TGV gate at each intersection of side and main channels.

The devices were designed using Autodesk Fusion (Autodesk, San Francisco, California). This software produced computer-aided design (CAD) and computer-aided machining (CAM) G-code (.simpl) files compatible with a Datron Neo computer numerical control (CNC) mill (Datron Dynamics, Milford, New Hampshire). Devices were milled on 3.175 mm poly(methyl)methacrylate plates (#8560K239; McMaster-Carr, Santa Fe Springs, California) for flow experiments with nonanol or 2.00 mm PS plates (#ST31-SH-000200; Goodfellow Corporation, Pittsburgh, Pennsylvania) for experiments with aqueous fluids. The channels were milled to have rounded inner corners to prevent the formation of capillary filaments using endmills with a cutter diameter of 1/32″ (TR-2-0312-BN) or 1/64″ (TR-2–0150-BN) from Performance Micro Tool (Janesville, Wisconsin). The accuracy of each device’s dimensions was verified using a Keyence wide area 3D measurement VR-5000 profilometer (Keyence Corporation of America, Itasca, Illinois). Channel floors in the devices were estimated to have a few microns of roughness due to the milling process. This surface roughness falls below the roughness values observed by Lade *et al.* to produce substantial fluctuations in velocities in capillary flow.^40^

Residues and debris were cleaned from the devices through ultrasonication in 70% (v/v) aqueous ethanol for 30 min using a Branson M2800H. The cleaning solvent was reused no more than five times. After sonication, the devices were rinsed in fresh 70% (v/v) aqueous ethanol and subsequently in deionized (DI) water. The devices were then left partially covered in a bioassay dish to dry in a fume hood overnight. Devices designed for aqueous solutions (PS devices) were oxygen-plasma treated with a Diener Zepto PC EX Type PB plasma treater and used within 5 minutes of plasma treatment.

Paper pads for the devices were designed in Adobe Illustrator 2023 (Adobe, San Jose, California) as 15.2 mm wide and 25 mm long rectangular pieces. This design was then cut out of Cytiva Whatman #1 filter papers (#1001-185) using a Graphtec CE-7000 plotter cutter and the Cutting Master 5 program (Graphtec America, Irvine, California). Paper pads were stored in a bioassay dish before use.

### Solvent preparation

For trials conducted on poly(methyl)methacrylate, dyed solutions of nonanol (Sigma-Aldrich, #131210) were made using Sudan I (Sigma-Aldrich, # 103624), Sudan III (Sigma-Aldrich, # S4131), Solvent Green 3 (Sigma-Aldrich, #211982), and Solvent Yellow 7 (#S4016), Oil Blue N (Sigma-Aldrich, #391557), Malachite Oxylate (Sigma-Aldrich, #M9015), and Methyl Violet (Sigma-Aldrich, #69710) each at concentrations of 0.5 mg mL^-1^. These solutions were originally made in 10 mL volumetric flasks at concentrations of 1 mg/mL and were subsequently diluted with nonanol to concentrations of 0.5 mg mL^-1^. For trials conducted on polystyrene, 0.6% dyed aqueous solutions were made using assorted food colorings (McCormick). These solutions were originally made in 10 mL volumetric flasks and later transferred to vials that were sealed for storage.

### Open-channel flow experiments

The devices were positioned on a white background atop an adjustable lab jack. A Nikon-D5300 ultrahigh-resolution single-lens reflective camera was used to capture the progression of solvent flow and fluid layer of each device at 60 fps. Tabletop photography lights were adjusted for each video to reduce shadows. Dyed nonanol was used to demonstrate fluid layering and better visualize flow.

For the devices with 0.7 mm step heights (Figure 3, 4), 71.6 µL of five to seven different dyed solutions were added to each side channel reservoir. For the device with 0.6 mm step height (Figure 3), 57.0 µL of dyed solution was added to each side channel reservoir. In both cases, the pinning at each gate was visually confirmed. Afterwards, 490 µL of dyed solution was added to the main reservoir. Devices were left to flow, and each TGV was released sequentially. For the travel distance and fluid velocity experiments, data collection stopped when the fluid reached the paper pad. For layering experiments, only two dyed solutions were used. These solutions were added to each reservoir in an alternating manner to enhance contrast between the layers for better visualization of the fluid layering. Data collection during layering experiments stopped after the resolution of layers through the camera was ensured.

**Figure 1.**
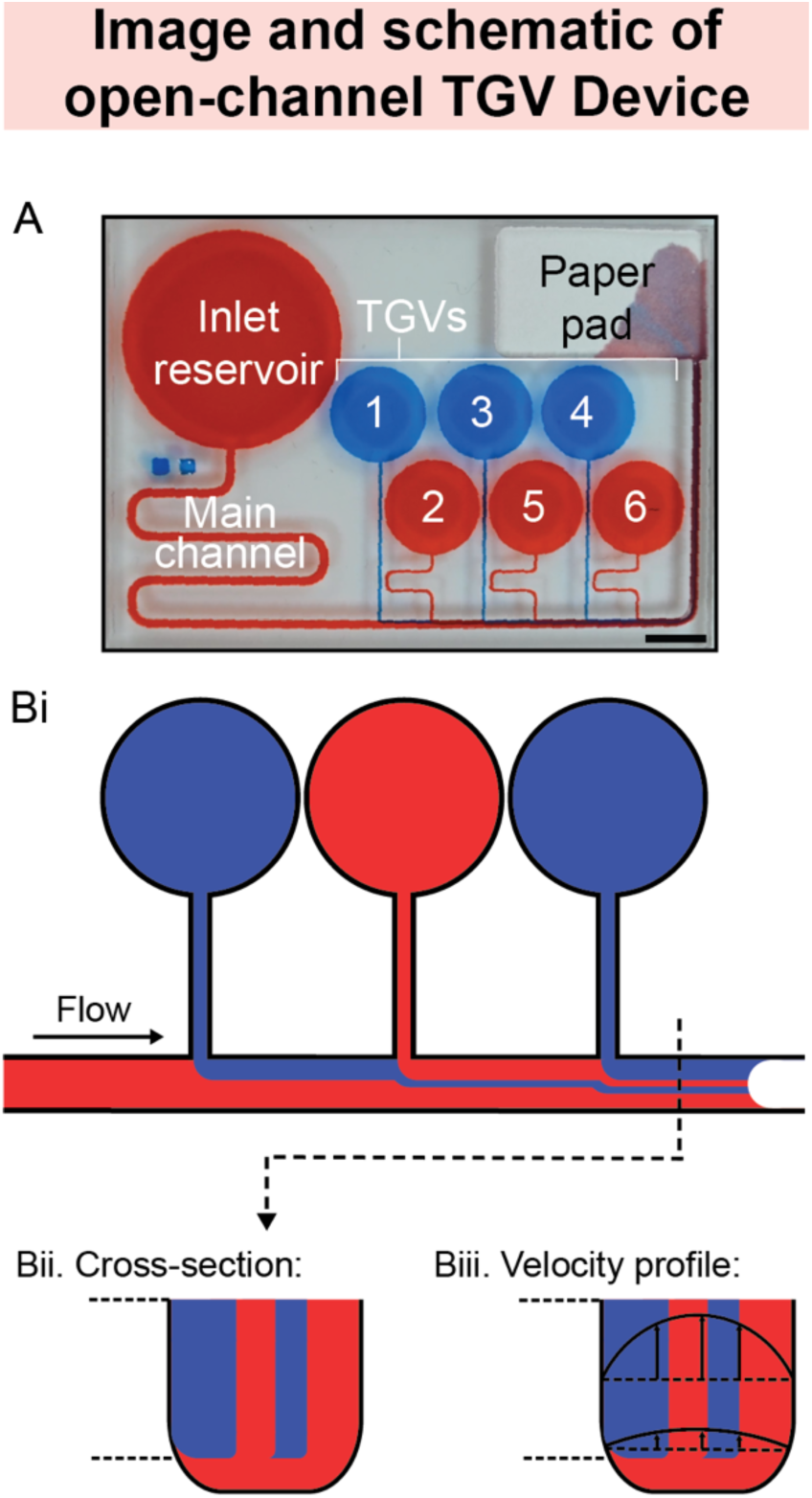
Annotated image and schematic of a TGV device after flow. (A) Image of an open-channel trigger valve device with six trigger valves. (Bi) A schematic representation illustrating the flow and layering of different miscible fluids in the main channel of TGV devices. (Bii): schematic view of the layers in a cross-section; (Biii): approximate Poiseuille velocity profiles in a cross-section.

**Figure 2.**
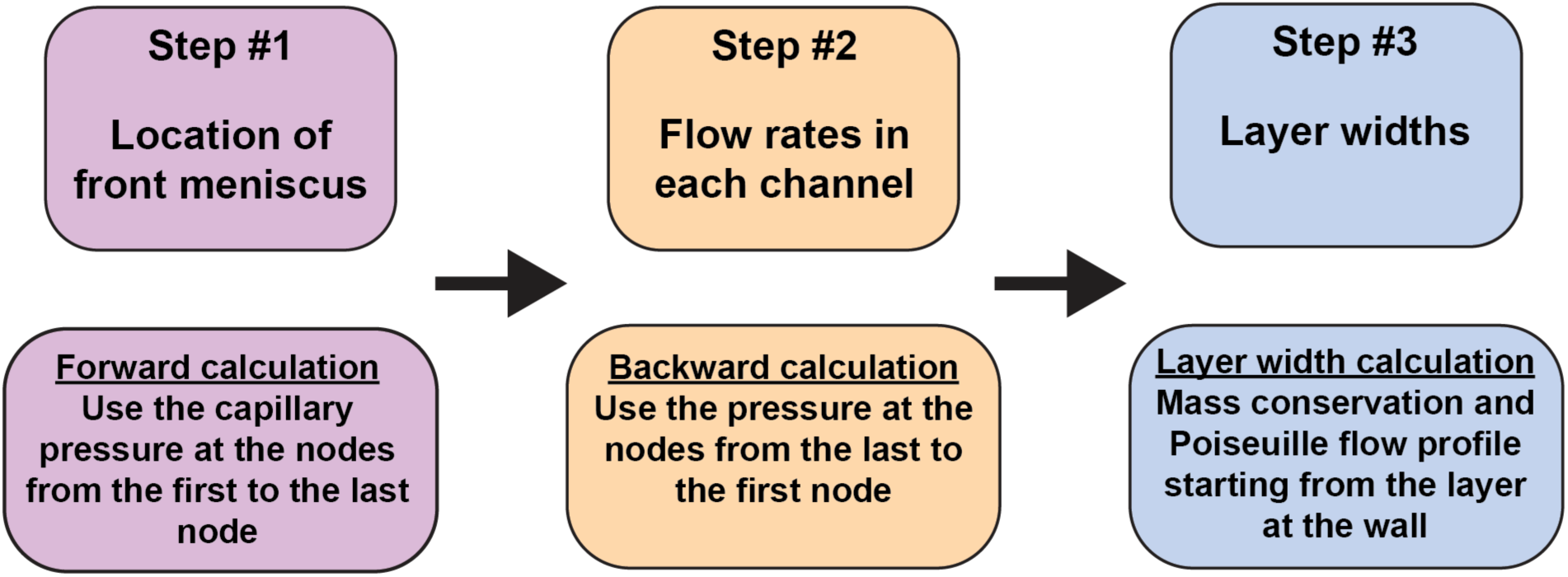
Overview of the three-step process used in the theoretical model to determine the layer widths created.

**Figure 3.**
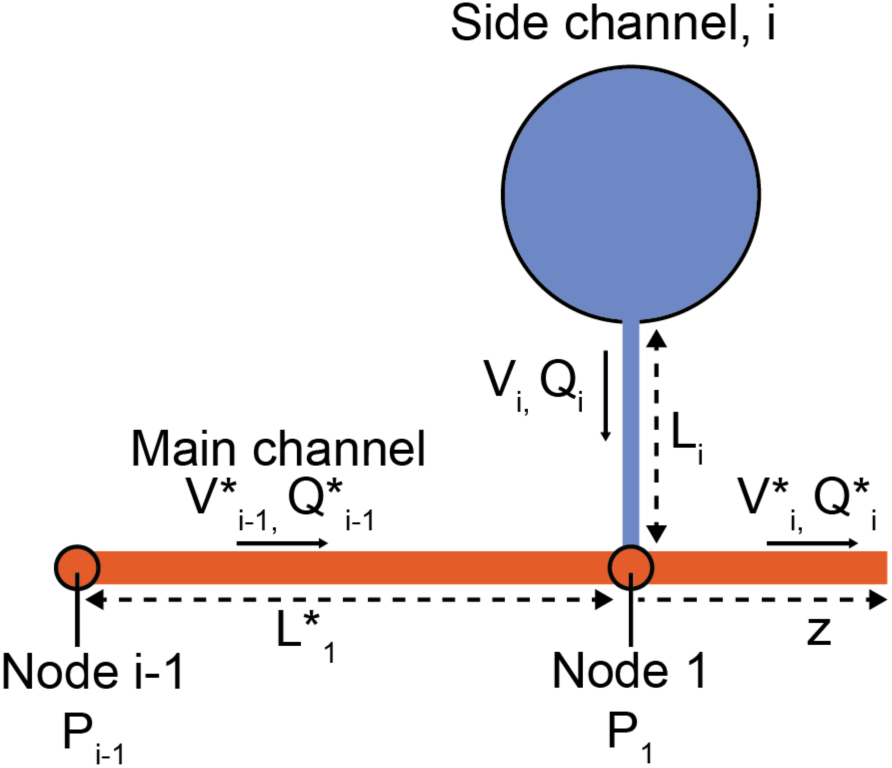
Sketch of main of TGV node I. The main channel is orange and side channel (#i) shown in blue. For the notations, an asterisk (*) always denotes the main channel. Here, Q stands for the flow rate and V js the velocity. This figure has been adapted from Tokihiro *et al*.^16^

**Figure 4.**
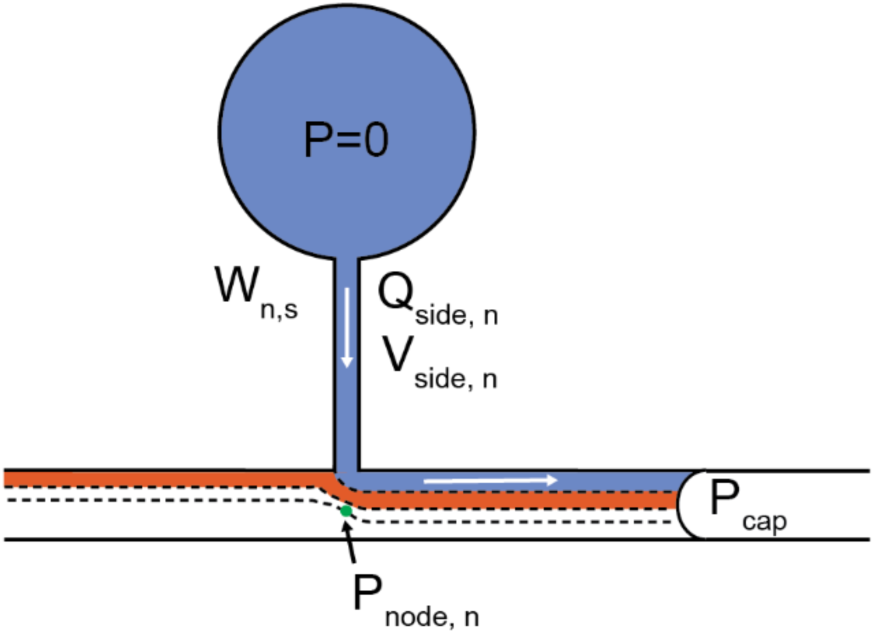
Principle for the calculation of the flow rate *Q̄_n_*

To analyze the recorded videos, a single image was extracted every 10 frames by a custom Python program^12^ designed for travel distance and velocity analysis. The position of the fluid front was tracked using the segmented line and measure tools in ImageJ (National Institutes of Health). The measurements were then exported as a .csv file.

The layer widths were determined 1 to 3 mm past each valve after the final valve of the device had been released. Measurements of “color” dyed fluids were analyzed using the segmented line and measure tools in ImageJ. These measurements were taken from a single frame for most layers. In some cases, a second frame after physically zooming the camera in was utilized for enhanced clarity of smaller layers. The measurements were then exported as a .csv file.

### Cell culture maintenance - 3T3 mouse fibroblasts

Cell patterning experiments were conducted with NIH/3T3 mouse fibroblast cells (ATCC). Cells were maintained in tissue culture flasks containing DMEM (Corning) supplemented with 10% fetal bovine serum (Gibco) and 1% penicillin-streptomycin (Gibco) at 37 °C, 5% CO_2_. Culture medium was replaced every 48 h until cells reached 90-95% confluency. To passage the cells, the culture was rinsed with 1X PBS (phosphate-buffered saline), followed by addition of TrypLE Express (Gibco). After incubation for 3 min at 37 °C, the TrypLE was inactivated by diluting with cell culture medium. The detached cells in liquid suspension were centrifuged at 300 RCF for 5 min. The cell pellet was resuspended in cell culture medium for further passaging or used for patterning experiments described below. 3T3 cells between passage numbers 10 and 15 were used for experiments.

### Patterning 3T3 cells in collagen with two-trigger valve device

Polystyrene trigger valve devices were plasma treated 30 to 60 minutes prior to cell patterning. The devices were treated with UV light for 15 minutes for sterilization. After the 3T3 cells were detached from the culture flask, the cells were dyed with CellTracker Green CMFDA or CellTracker Red CMTPX at a final concentration of 25 µM in DMEM for 30 min at 37 °C. After the dyeing process, the cells were pelleted and resuspended at a final concentration of 2 to 3×10^6^ cells mL^-1^ in a collagen extracellular matrix precursor solution. The collagen solution was prepared by mixing a stock of rat tail collagen type I (Corning) in 0.02 N acetic acid with sterile deionized water, 10X PBS, 1 M HEPES buffer solution, 7.5% sodium bicarbonate solution, and 1 N NaOH to achieve a final collagen density of 3 mg mL^-1^ and kept on ice to prevent gelation. 100 µL of cell-laden liquid collagen solution was pipetted into each of the trigger valve channel reservoirs, and 630 µL of cell-laden liquid collagen solution was pipetted into the main channel reservoir. After the fluid front of the main channel reached the second trigger valve, the device was incubated at 37°C for 30 min to allow for collagen gelation. Devices were then sealed with microplate sealing film and imaged with 10X and 20X magnification on a Zeiss LSM 900 confocal microscope equipped with ZEN software.

### Theory

In this work, we use a series of trigger valves (TGV) to form co-flowing layers that pile up after each TGV release as schematically shown in Figure 1.

The algorithm developed in Tokihiro *et al*.^16^ produces the location of the advancing front meniscus in function of time. It is the Step #1 of the numerical program. We have further developed the algorithm to account for the flow rates in the different channels in function of time (Step #2) and the layer width of the different laminated flows in the main channel (Step #3).

It is recalled that step #1 calculation—provided here for ease of reference and detailed in Tokihiro *et al.*^16^ —progresses from the first node (intersection of the main channel with the first side channel) towards the last node (intersection of the main channel with the last side channels). The velocity of the meniscus is determined using the mass conservation equation at each node, the pressure drop between each node, and the capillary pressure at the front meniscus. We present here the two next steps; calculating the flow rates in each channel and finally the layer widths of each additional fluid. A detailed table of notations and the corresponding definitions can be found in Table SI.4.1.

#### a. Step #1

The following section (Step #1) has been adapted from Tokihiro *et al.* and is reproduced here for ease of reference.^16^ The position of the front meniscus is determined by calculating its travel distance along the main channel. Each junction between a side channel and the main channel is defined as a node. The calculation proceeds recursively from one node to the next. Figure 3 illustrates the (i)-th trigger valve gate (TGV).

The pressure P_i_ can be expressed by two linear relations associated with two segments (main channel #i-1 and side channel #i)

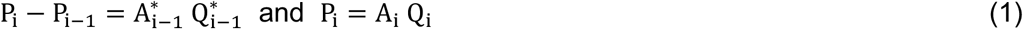

where A_i_ and - A^∗^are constants linked to the channel and liquid characteristics. P_i-1_ is known from the previous step (recurrence). Combining these two relations one finds the relation between Q_i_^∗^and P_i_ or V_i_^∗^
and P_i_

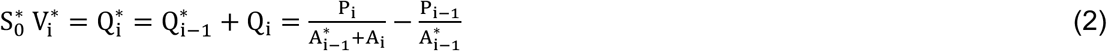

Now, considering the portion of main channel #i, considering the capillary pressure P_cap_, we can write

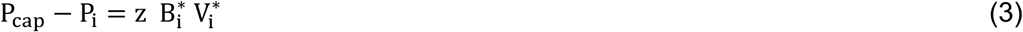

where B_i_^∗^ is a constant depending on the channel and liquid characteristics. Using V_i_^∗^ = dz⁄d t, combining (2) and (3) yields the quadratic equation for the travel distance z

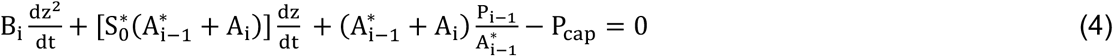

The capillary pressure P_cap_ is a constant due to the uniformity of the main channel (and to the viscous regime in which the constant angle is constant). The constants in equation (4) are 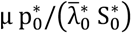, where p^∗^ is the cross-section perimeter of the main channel (constant from inlet to outlet) and λ̄^∗^ the friction length, 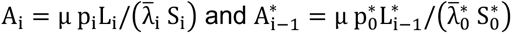:.

#### b. Step #2

The flow rates in the system are calculated using a backward approach (Figure 4): Considering the last node *n*, the pressure *_Pn_* at the node *n* is given by

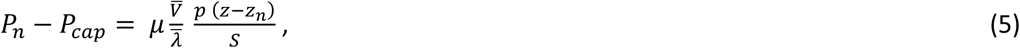

where *p* is the total perimeter of a cross-section, *S* the cross-sectional area of the main channel, z the axial coordinate of the location of the meniscus, *z_n_* the location of the node *n*. As noted in Tokihiro *et al.*^16^, *μ* is the liquid viscosity, *V̄* the average velocity in the main channel, λ̄ the average friction length (λ̄*=1/3 *(w+h)/(w/h+2h/w))*, and *_Pcap_* the capillary pressure at the meniscus.

At a distance *z*, as *_V̄_* is known from step #1, relation (5) produces the pressure at the node *_Pn_*. The mean velocity *V̄_n_* in the side channel *n* is then deduced from *_Pn_*, where *L_n_* is the length of the side channel,

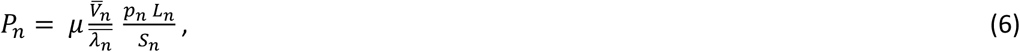

and the flow rate in the side channel *n Q̄_n_* is

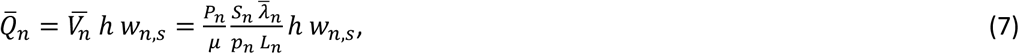

where *w_n,s_* is the width of the side channel n. Using the mass conservation equation, the flow rate in the main channel between nodes *n-1* and *n* denoted 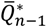 is then

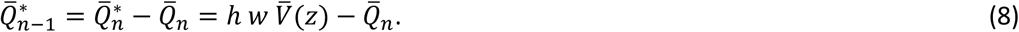

Note that the asterisk denotes the main channel. The scheme can be reproduced to the rank *n-1* to determine the pressure at any given node.

#### c. Step #3

After each TGV, a new layer forms. Therefore, at node j, there are the j layers corresponding to the j upstream TGVs and the layer corresponding the main flow, for a total of j+1 layers. Without loss of generality, we present the layer width calculation after the last (index n) TGV. To determine the layer widths, first consider the layer next to the wall (wall where the TGVs are placed). Let *n* be the index for this side channel and layer (Figure 5).

**Figure 5.**
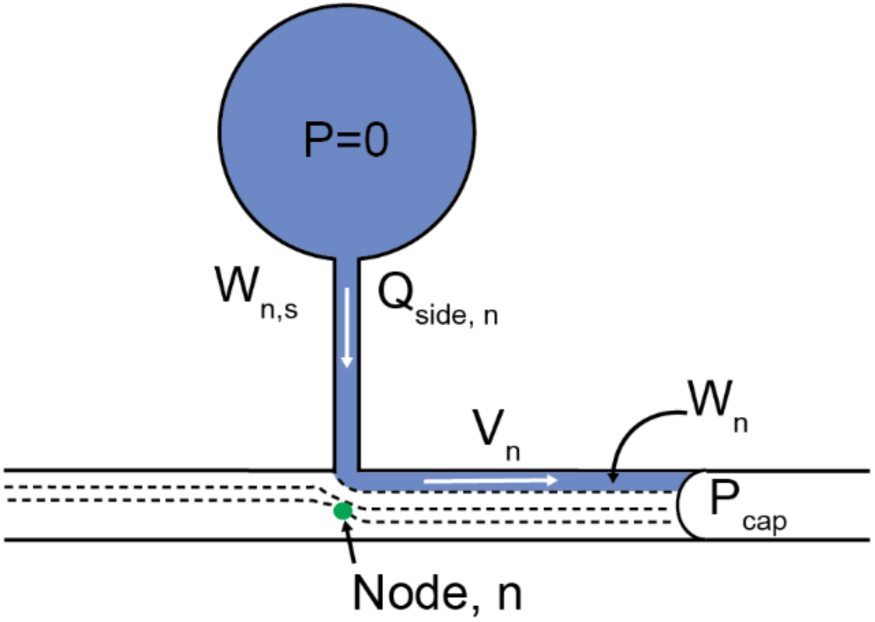
Calculation of the layer width wn: as there is no noticeable mixing, the flow rate in the layer is equal to that flowing in the side channel *n*.

### Mass conservation equation yields

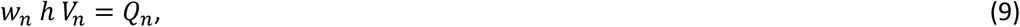

where *h* is the layer depth—approximately the depth of the side channels due the negligible diffusion at the bottom—and *V_n_* the average velocity in the n^th^ layer which we assume to be the same as the velocity at the middle of the layer, because of the very small layer width associated to the number of TGVs. Assuming that the velocity profile in the main channel is a Poiseuille profile, the velocity in a cross section is given by

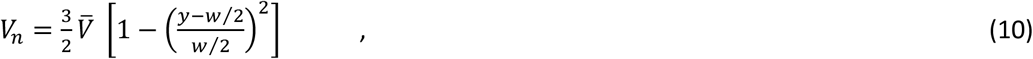

where *w* is the main channel width, *V̄* the average velocity in the main channel—calculated in step #1—and *y* the coordinate perpendicular to the wall (*y*=0 at the wall). In the case of many small layers, the average velocity of the layer is approximated by the velocity at the layer center. If *y_1,n_* and *y_2,n_* are respectively the coordinates of the layer boundaries (at the wall *y_1_*=0), the velocity of the layer is

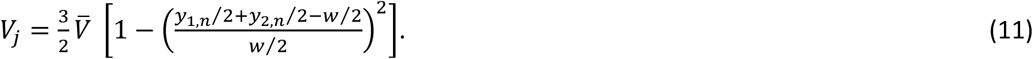

Remarking that

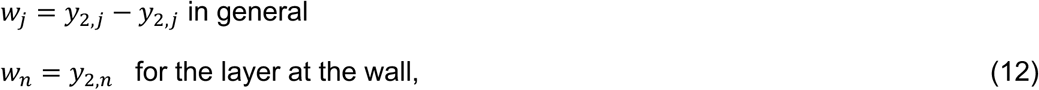

substitution of (11) and (12) in (10) first produces an equation in *w_m_* (root of a cubic polynomial). It was found easier to use an iterative method. The first approximate of *w_n_* denoted *w*^(1)^is

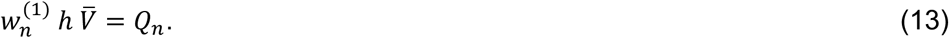

Then *V_n_*^(1)^is obtained using (11) and re-injected in (9) to obtain a new *w_n_*^(2)^, and so on until convergence. The scheme is repeated for each layer from *n* to *1* to determine the widths of each layer. The remaining layer width *w_0_* corresponding to the main flow is obtained by difference *w*_0_ = 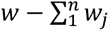.

## Results and Discussion

As shown in the previous section we introduce a numerical model for predicting the widths of coflowing fluid layers within an open capillary TGV device. Here we provide a comparison to experimental data using devices micromilled in PMMA plastic and nonanol as the flowing liquid. In this work, we have investigated the effect of fluid layering through manipulations of varying step heights and the number of trigger valves. We show experiments conducted in high viscosity liquids such as nonanol and 0.65% m/m glycerol-water solutions to ensure distinct coflowing layers through the end of the channel and also serve as a model for high-viscosity liquids such as hydrogel precursor solutions for biologically-relevant applications. Furthermore, nonanol and aqueous glycerol solutions also have suitable contact angles to achieve spontaneous capillary flow in PMMA and PS devices. Here, we also show the versatility of these devices utilizing aqueous fluids in PS devices with a proof-of-concept application using 3T3 fibroblasts in a collagen extracellular matrix.

Trigger valve devices used in this study have a large inlet reservoir designed to have consistent Laplace pressure. This reservoir allows fluid to flow into a channel where flow is initially in an inertial flow regime but slows into a viscous flow regime before arriving at a side channel opening. As the fluid flows past these side channels the TGVs are released allowing for the side channel fluid to flow alongside the main channel flow until eventually reaching an end reservoir holding a paper pad (Figure 1A). Fluid from each TGV flows in the same direction through the main channel and maintains a distinct layer until reaching the paper pad at the end of the channel. In this work a model has been developed to predict these layer widths and fluid contributions to the main channel flow.

### Fluid layering and flow in devices with different step heights

A critical parameter in open-channel trigger valve systems is the step height–the distance between the floor of the TGV channel and the main channel floor. In our previous work, we investigated TGVs with a 0.4 mm step height (0.6 mm deep TGVs) and have now been able to implement in this work both 0.6 mm and 0.7 mm TGV channels. In this section, two devices containing five trigger valves were designed and manufactured. The only difference between devices was the step height of each TGV. One device contained TGVs with depths of 0.6 mm, corresponding to a 0.4 mm step height. The second device had a 0.7 mm TGV depth, leading to a 0.3 mm step height. Dyed nonanol solutions were then flowed through each device so that the widths of each layer could be recorded and measured experimentally (Figure 5A). Figure 5A shows the top-down view of the device with a 0.6 mm side channel depth after the last trigger valve was released (Figure 6Ai) along with a profilometry image of the intersection of the trigger valve and main channel (Figure 6Aii). Figure 5Aiii shows images of the main channel where the layer width measurements were taken (∼3 mm downstream of each TGV). A corresponding set of images for a device with a 0.7 mm side channel depth can be found in Figure SI.2.3. These results were then compared to the theoretical model for validation (Figure 6B and C). For the device with 0.6 mm depths, the layer widths after release of the final valve averaged over three trials were 0.356 mm for the main fluid, 0.257 mm, 0.128 mm, 0.102 mm, 0.0825 mm, and 0.064 mm for TGVs five to one (Figure 6B). For the device with 0.7 mm depths, the layer widths after release of the final valve averaged over three trials were 0.330 mm for the main fluid, 0.248 mm, 0.133 mm, 0.113 mm, 0.095 mm, and 0.075 mm for valves five to one (Figure 6C). This corresponds well to the theoretical trend that earlier layer widths continually decrease as further TGVs are released. These values also align well with the predicted layer widths and are an average of ∼7 percent different when there are 0.3 mm step heights and ∼12 percent different when there is a 0.4 mm step height. Average and standard deviation calculations can be found in Tables SI.3.1 and SI.3.2.

**Figure 6.**
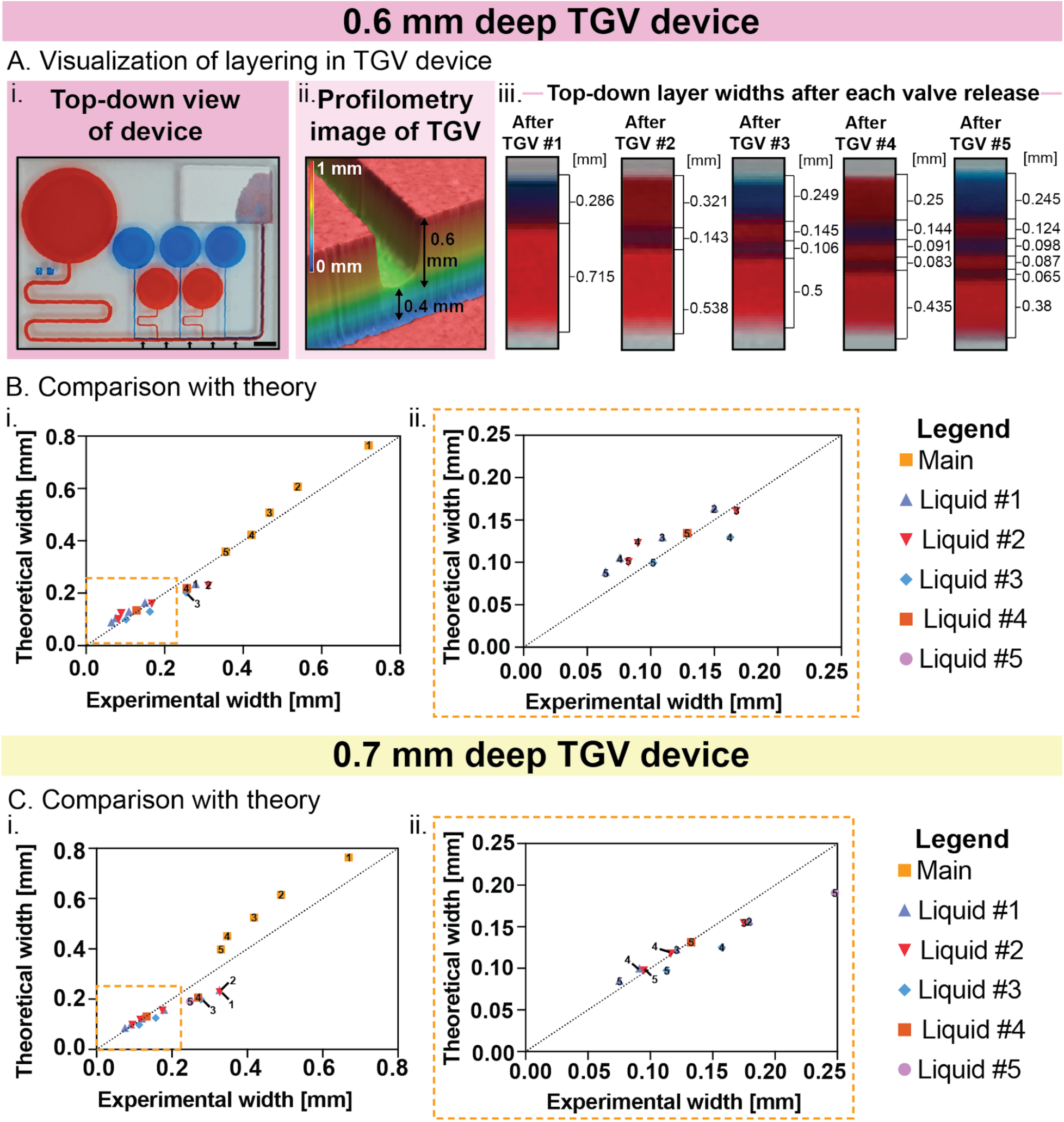
Layering in TGV devices with varying step heights. (A) i. Top-down image of fluid layering after the release of the final TGV in a 0.6 mm deep TGV device. Arrows correspond to the position in the channel that layer widths are measured following TGV release. The scale bar is 5 mm. ii. Profilometry image of a 0.6 mm deep TGV entrance into the main channel. iii. Images of fluid layering in the main channel after each TGV release for a 0.6 mm deep TGV device. (B) Comparison of experimental layer width after the release of the final TGV in all parts of the channel to the theoretically predicted width (i), including a zoomed-in graph of the boxed region (ii). Colored shapes correspond to liquid released from its respective TGV, and numbers within the shapes correspond to the liquid’s width immediately after the release of that numbered TGV. For example, the width of Liquid #4 is shown immediately after the release of TGV 4 and additionally after TGV 5. Data is shown as the average of 4 individual devices (n = 4). (C) Comparison of experimental layer width (i) after the release of the final TGV in all parts of the channel to the theoretically predicted width, including a zoomed-in graph of the boxed region (ii). Colored shapes correspond to liquid released from its respective TGV, and numbers within the shapes correspond to the liquid’s width immediately after the release of that numbered TGV. Following previous logic, the width of Liquid #4 is shown immediately after the release of TGV 4 and additionally after TGV 5. Data is shown as the average of 4 individual devices (n = 4). Layer widths for the individual trials are provided in Tables SI.3.1 and SI.3.2.

The theoretical model used in this study also has the capability to predict the travel distance (and therefore the velocity) of fluid in TGV systems. This capability had previously been validated for devices containing three TGVs, all with depths of 0.6 mm. To further test the model, dyed nonanol was flowed through the devices to measure the movement of the fluid front. The experimental data (average of four trials in a device with a 0.6 mm depth) correlate closely with the theoretical model (Figure 7A, C). This is also seen after an average of four trials in the device with 0.7 mm depths (Figure 7B, D).

**Figure 7.**
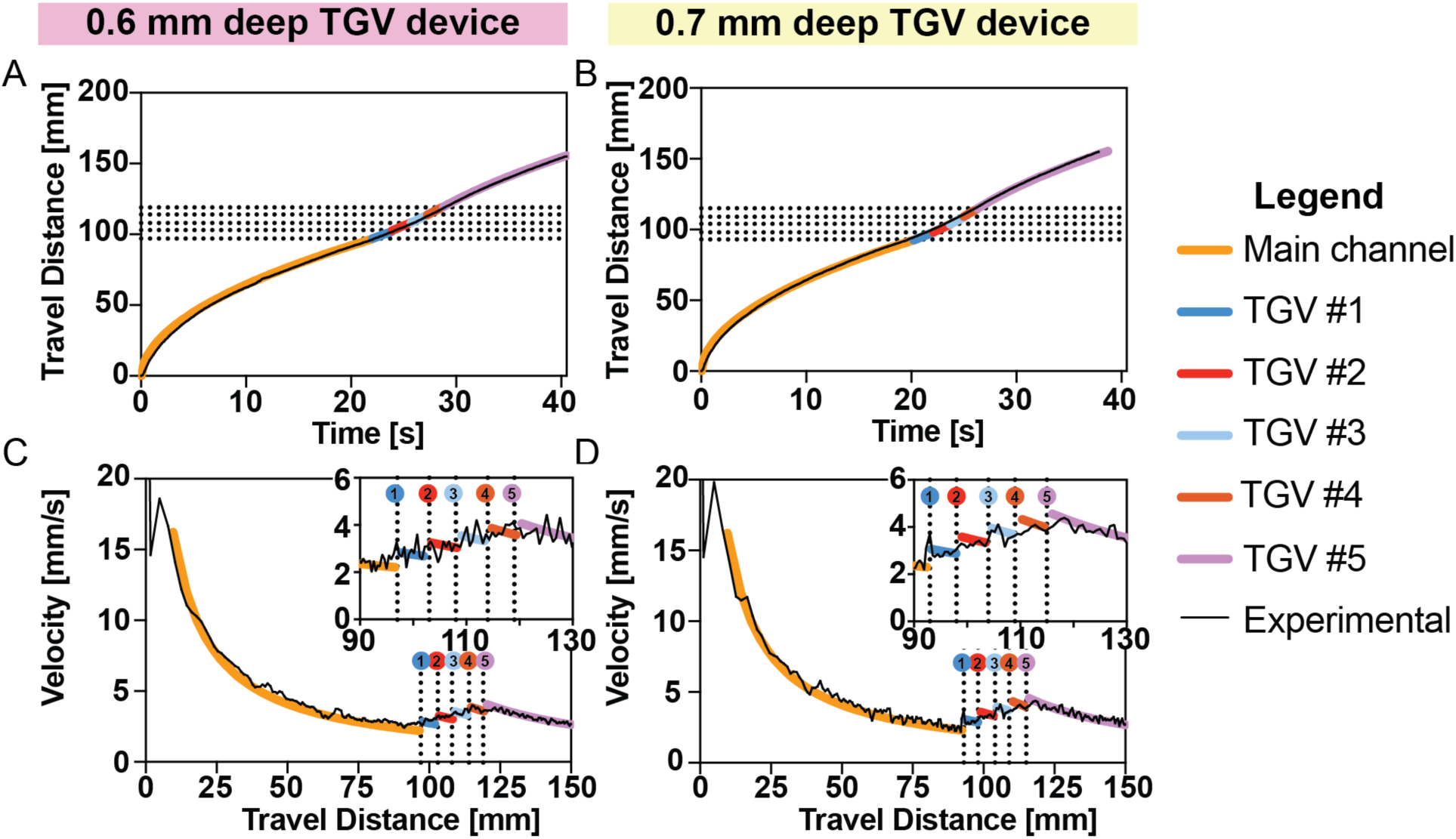
Flow in TGV devices with varying step heights (A) Travel distance over time in devices with 0.6 mm deep TGVs (B) Travel distance over time in devices with 0.7 mm deep TGVs (C) Velocity over travel distance in devices with 0.6 mm deep TGVs (D) Velocity over travel distance in devices with 0.7 mm deep TGVs. A representative trial is shown for all plots shown. Data for n = 4 can be found in Figures SI.2.1 and SI.2.2 for the devices with 0.6 mm and 0.7 mm deep TGVs, respectively.

### Fluid layering and flow with up to 7 trigger valves

Increasing the number of TGVs in a device increases the potential use in multi-step reactions and overall control of fluid behavior. Therefore, we desired to expand and validate the theoretical model in cases with larger amounts of trigger valves. To this end, we designed and manufactured devices containing six and seven trigger valves. In both designs, all TGVs had a depth of 0.7 mm (0.3 mm step heights), causing the only difference in devices to be the number of TGVs. Dyed nonanol solutions were flowed through each device so that the widths of each layer could be recorded and measured experimentally (Figure 8A,C). These results were then compared to the model for validation (Figure 8B, D). Figure 8Ai and 8Ci shows the top-down view of after the last trigger valve was released for 6-valve and 7-valve devices, respectively. Figure 8Aii and 8Cii show images of the main channel where the layer width measurements were taken (∼3 mm downstream of each TGV). For the device with six trigger valves, the layer widths after the release of the final valve averaged over three trials were 0.327 mm for the main fluid, 0.243 mm, 0.128 mm, 0.081 mm, 0.081 mm, 0.073 mm, and 0.059 mm for valves six to one (Figure 8B). For the device with seven trigger valves, the layer widths after the release of the final valve were 0.292 mm for the main channel, 0.243 mm, 0.125 mm, 0.102 mm, 0.073 mm, 0.060 mm, 0.064 mm, and 0.049 mm for valve seven to one (Figure 8D). Comparing these results to the model’s predicted layer width gives an average difference of 12% for the device with six TGVs and 8.5% for the device with seven TGVs, showing that the model can predict the layer width of devices with increased valves well. Average and standard deviation calculations can be found in Tables SI.3.3 and SI.3.4.

**Figure 8.**
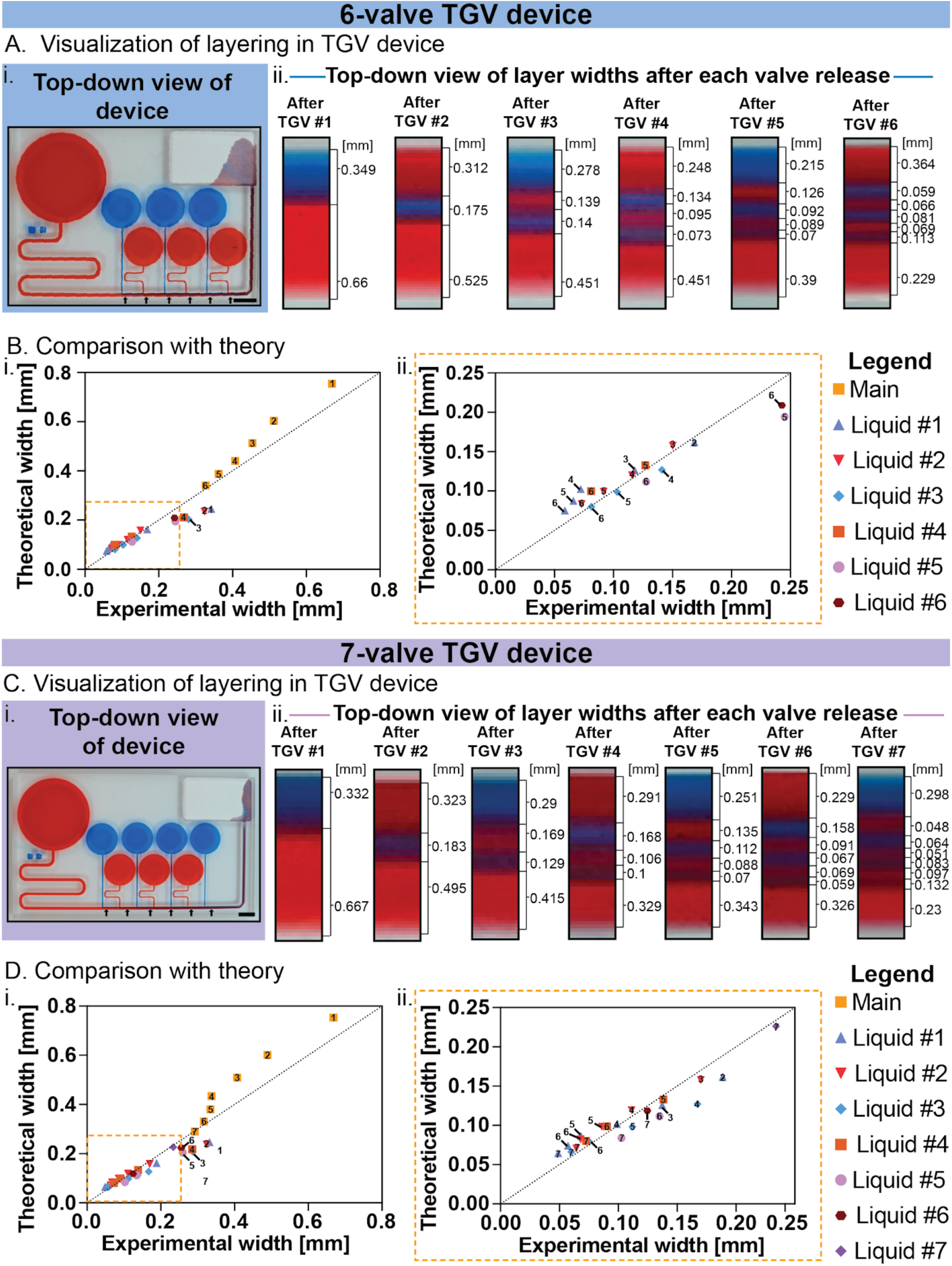
Layering and flow in devices with six and seven TGVs. (A) i. Top-down image of fluid layering after the release of the final TGV in a 6 TGV device. Arrows correspond to the position in the channel that layer widths are measured following TGV release. Scale bar is 5 mm. ii. Images of fluid layering in the main channel after each TGV release for a 6 TGV device. (B) Comparison of experimental layer width after the release of the final TGV in all parts of the channel to the theoretically predicted width (i), including a zoomed-in graph of the boxed region (ii). Colored shapes correspond to liquid released from its respective TGV, and numbers within the shapes correspond to the liquid’s width immediately after the release of that numbered TGV. For example, the width of Liquid #4 is shown immediately after the release of TGV 4 and additionally after TGV 5 and 6. Data is shown as the average of 4 individual devices (n = 4) (C) i. Top-down image of fluid layering after the release of the final TGV in a 7 TGV device. Scale bar is 5 mm. ii. Images of fluid layering in the main channel after each TGV release for a 7 TGV device. (D) Comparison of experimental layer width (i) after the release of the final TGV in all parts of the channel to the theoretically predicted width, including a zoomed-in graph of the boxed region (ii). Colored shapes correspond to liquid released from its respective TGV, and numbers within the shapes correspond to the liquid’s width immediately after the release of that numbered TGV. Following previous logic, the width of Liquid #4 is shown immediately after the release of TGV 4 and additionally after TGV 5, 6, and 7. Data is shown as the average of 3 individual devices (n = 3). Layer widths for the individual trials are shown in Tables SI.3.3 and SI.3.4.

Along with the validation of layer widths, we validated the travel distance and velocity of the main fluid front predictions of the expanded model. To do this, dyed nonanol solutions were flowed through the devices, and the fluid front meniscus position was measured from the inlet reservoir to the outlet immediately before contact with the paper pad. Experimental travel distance and velocity measurements of the six and seven TGV devices for representative trials were compared to the model (Figure SI.2.3). Raw data plots for n = 3 for both the 6-valve and 7-valve devices can be found in Figures SI.2.4 and SI.2.5, respectively. These results correspond well to the model’s prediction, showing its expanded capability for larger TGV devices.

### Validation of theoretical model with aqueous solutions and proof-of-concept application of open trigger valve system with 3T3 cells in a collagen extracellular matrix

To show greater diversity in the application of TGV systems and validate our model under different conditions, we created a device suitable for aqueous solutions. To this end, we changed the plastic being used from native PMMA to oxygen-plasma treated PS, a plastic commonly used for biological applications. The contact angle of water on oxygen-plasma treated PS has a lower contact angle when compared to native PMMA. This allows for similarly designed TGV devices to those used with nonanol to be used with aqueous solutions. Keeping the device design the same as what was previously used for PMMA devices in Figure 6C, glycerol-water solutions (0.65% m/m) dyed with food coloring were flowed through the devices to measure the movement of the fluid front. The travel distance and velocity results measured through this flow were then compared to the theoretical model for validation (Figure 9A). Through this comparison the experimental data closely correspond with the theoretical model. For further validation of the expended model, the layer widths of the aqueous solution flow were measured. For this device and fluid combination, the layer widths after release of the final valve averaged over three trials were 0.384 mm for the main channel fluid, 0.078 mm, 0.89 mm, 0.091 mm, 0.136 mm, and 0.198 mm for valves one through five (Figure 9B, C). Comparing these widths to the theoretical values result in an average of less than one percent difference. These results are critically important as compatibility with PS can expand the application of these devices as PS is extensively used in cell culture platforms.

**Figure 9.**
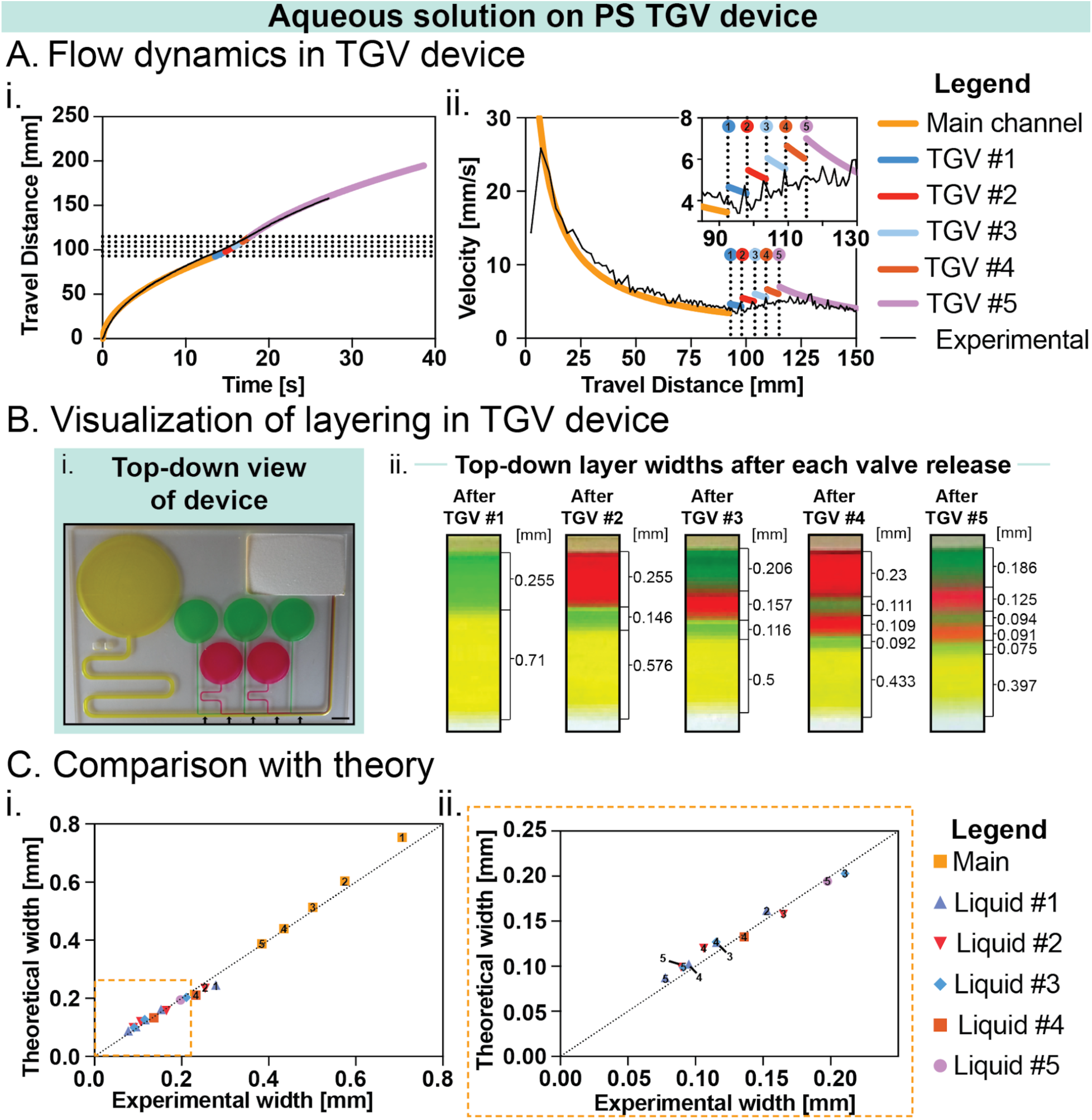
Flow and fluid layering of aqueous solution on oxygen-plasma treated PS 5-valve TGV device. (A) Flow dynamics in a 5-TGV oxygen-plasma treated PS device showing the distance traveled over time (i) and the calculated velocity along the main channel length (ii) for a representative trial. Travel distance data for n = 4 can be found in Figure SI.2.6. (B) i. Top-down image of fluid layering after the release of the final TGV in the 5-valve TGV device. Arrows correspond to the position in the channel that layer widths are measured following TGV release. The scale bar is 5 mm. ii. Images of fluid layering in the main channel after each TGV release for a 6 TGV device. (B) Comparison of experimental layer width after the release of the final TGV in all parts of the channel to the theoretically predicted width (i), including a zoomed-in graph of the boxed region (ii). Colored shapes correspond to liquid released from its respective TGV, and numbers within the shapes correspond to the liquid’s width immediately after the release of that numbered TGV. For example, the width of Liquid #4 is shown immediately after the release of TGV 4 and additionally after TGV 5 and 6. (C) i. Top-down image of fluid layering after the release of the final TGV in a 7 TGV device. ii. Images of fluid layering in the main channel after each TGV release for a 7 TGV device. (D) Comparison of experimental layer width (i) after the release of the final TGV in all parts of the channel to the theoretically predicted width, including a zoomed-in graph of the boxed region (ii). Colored shapes correspond to liquid released from its respective TGV, and numbers within the shapes correspond to the liquid’s width immediately after the release of that numbered TGV. Following previous logic, the width of Liquid #4 is shown immediately after the release of TGV 4 and additionally after TGV 5. Data is shown as the average of 3 individual devices (n = 3). Layer widths for the individual trials are shown in Table SI.3.5.

To demonstrate applicability of this device for a broader audience such as those interested in organ-on-a-chip or hydrogel patterning in a three-dimensional extracellular matrix (ECM), we used our open capillary trigger valve system to layer cells in a collagen extracellular matrix. The devices were fabricated similarly—in oxygen-plasma treated PS—to our previous device that used aqueous solutions as the collagen ECM is largely aqueous. Two new considerations were taken during device design phase: (1) the extension of the channel length after the last trigger valve to preserve the layers and keep the fluid in the main channel during the gelation process and (2) preserving a small footprint for imaging on the confocal microscope system. An engineering drawing of the device can be found in Figure SI.1.2. In a 2 TGV system, we used red and green fluorescent 3T3 mouse fibroblasts laden in a 3 mg mL^-1^ collagen ECM gel precursor solution for this proof-of-concept experiment (Figure 10A). After the collagen ECM precursor solidified, we used confocal fluorescence microscopy, we were able to image a region after the trigger valves were released and the collagen was incubated to gel. A maximum intensity projection image of the z-stacked layers is shown in Figure 10B with distinct red and green regions demonstrating the capacity of the device to layer biologically-relevant solutions with cells thereby creating a multi-region construct and allow for imaging on a microscope. Here, the 3T3 cell line was chosen for its well-characterized nature and robustness, but other cell lines that are used for layering or modeling adjacent regions of tissues could also be used. The orthogonal projection on the right side of the fluorescent image depicts the YZ side view of the image and shows minimal overlap or mixing at the top of the flow (in the z direction; near the air-liquid interface). However, there is some overlap of the red and green signal (shown in yellow) in the region that is closest to the floor of the channel due to the step in the z direction which is expected in stair-step trigger valves. This effect could be mitigated through the suspension of the device along two rails and allowing the trigger valves and main channel to be at the same channel depth—effectively removing the stair-step and the Z axis layering—which will be explored in future work. In addition, an accurate layer width measurement requires sophisticated image processing and increasing the cell concentration are other areas for further investigation. Nonetheless, we show here a method for hydrogel patterning parallel regions of cells in a pump-free method that also allows for direct access to the cells in the device.

**Figure 10.**
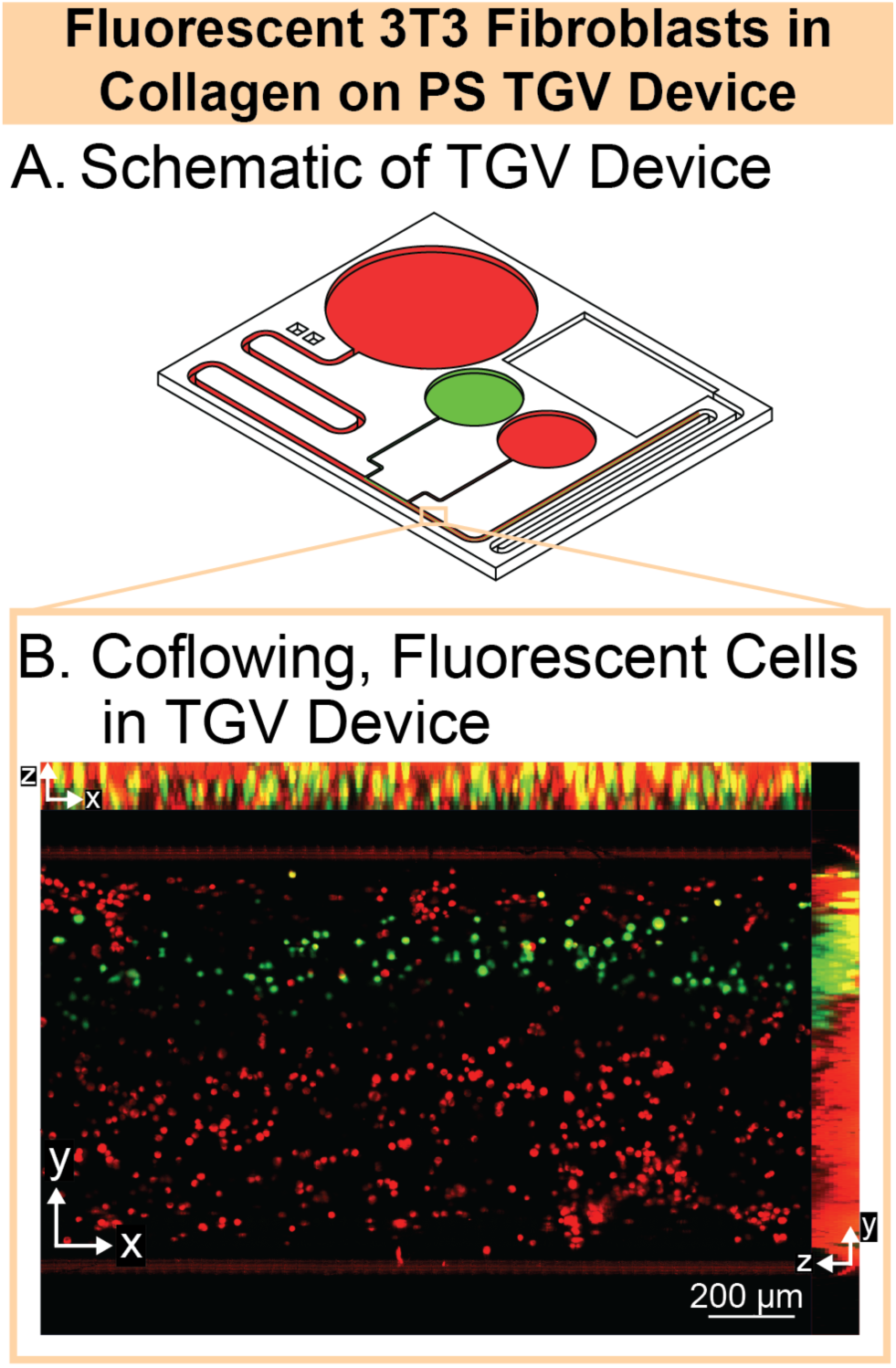
Coflow of red and green dyed 3T3 fibroblasts in collagen extracellular matrix gel precursor solution using a trigger valve device. (A) Schematic of a trigger valve device with two trigger valves and a longer main channel post-trigger valves for gelling of the extracellular matrix. (B) Representative maximum intensity projection image from bottom-up view (XY) of the main channel after the last trigger valve was released with layering of red and green fluorescent 3T3 fibroblasts. Orthogonal projections of the layers are shown on the top (XZ) and on the right (YZ) showing the z stacking of the layers. Experiment was done in three technical replicates (n=3). Images of all trials shown in Figure SI.5.1.

### Limitations of the present work and comparison with previously reported passive valves

In this study, we extended our previous work on implementation of trigger valves in open systems through the development of a theoretical model and comparison with in-lab measurements of the widths of each coflowing layer. One limitation of this work is that we have focused on the flow of high viscosity liquids such as nonanol, glycerol-water solutions, and hydrogel precursor solutions, but have yet to validate this with liquids of lower viscosity such as pure water or isopropyl alcohol. The main advantage of using viscous fluids for this demonstration is their low interlayer diffusion coefficients, which help to preserve the fluid layers over longer distances along the device. Another limitation of this work is that due to the passiveness of the device, the flow is unable to be halted mid-flow for dosing at specific time points and is generally a one-shot flow mechanism. Lastly, we are able to approximate flow rate of each layer through our theoretical model, but one effect seen in the fluorescent image (Figure 10B) is the overlap of the coflows at the bottom of the channel due to the stair-step nature of the trigger valves limiting our approximation. Elimination of the stair-step and more precise measurement of the flow rate are areas of future investigation.

We note that the contact angle will differ depending on the liquid and substrate chosen, so it will be important to consider the effect of varied contact angles to evaluate the generalizability of this method. We here provide a framework for open channels, but this method could be readily extended to closed channels by adjusting two parameters: the generalized Cassie angle (denoted by θ*) and the average friction length (denoted by ̅̄). These parameters are generally easier to determine in closed channels whose walls are made of the same material and produced using the same fabrication process. In this case, the generalized Cassie angle reduces to the conventional contact angle, (θ), while the average friction length for a closed channel with a rectangular cross-section can be estimated using a series-expansion approximation of the flow profile (see Bahrami *et al.*^41^ for an example). For closed channels with non-monolithic walls, the same approach as that used for open channels must be followed: the free surface is replaced by a solid wall, whose wettability and frictional properties must be taken into account.

In general, trigger valves are just one of many different passive valves that can control fluid flow via geometrical features and can be implemented in open channels. In this work, we focus on two-level, stair-step trigger valves which are one of the most stable capillary valves. While there is little in the way of valving in open channels in the literature, valving in closed channels—specifically, passive valves—are well-documented. Valving mechanisms rely on either channel geometry, channel wettability, or pressure changes. Some types of valves include capillary stop valves, TGVs, capillary delay valves, capillary retention valves, or siphon valves.^13,42–47^ Each confer their own strengths and limitations and this has been reviewed by others in the field.^48,49^ Choosing the valve type for a particular device ultimately depends on the use or application of the microfluidic device.

## Conclusion

This work expands the capabilities of autonomous fluid control in open microfluidic systems through the integration of advanced trigger valve designs. By showing the use of varied step heights, increasing the amount of trigger valves up to seven, demonstrating compatibility with fluids of differing properties, and utilizing varied plastics we broaden the functional range of trigger-valve-based open microfluidic platforms. We also show here a proof-of-concept application of layered patterning of 3T3 fibroblasts in an open TGV device through the flow of fluorescent fibroblast-laden collagen ECM precursor solution. The further development of a theoretical framework to predict fluid travel distance, velocity and layering width strengthens the reliability and design capabilities of these systems. Through focus on the layers produced through sequential TGV release we show stable and reproducible layers with widths as low at 50 microns. These layers were accurately determined as a function of the geometric characteristics of the different side channels of each trigger valve. These advances establish a foundation for accessible trigger valve device design and allow for more sophisticated devices to be created. Additionally, the open nature of these devices provides unique advantages to this framework over other techniques to generate layered co-flows. Being an open microfluidic system allows for the device to be made by diverse fabrication methods, to have greater accessibility to the sample, and simpler fluid control when compared to closed techniques. These devices would support applications in many research areas including at-home sample preparation, biosensing platforms, and especially hydrogel patterning of 3D cell culture.

## Acknowledgements

Research reported in this publication was supported by the National Institutes of Health National Institute of General Medical Sciences Grant R35 GM128648 (A.B.T.) and National Institute of Environmental Health Sciences K12 ES033584 (T.M.N.). We also acknowledge the M.J. Murdock Diagnostics Foundry for Translational Research. The content is solely the responsibility of the authors and does not necessarily represent the official views of the National Institutes of Health.

## Author Contributions

J.C.T., T.J.C., E.B., A.B.T., J.B. contributed to conceptualized the study design. J.C.T., T.J.C., A.L., J.M.W., X.S., I.H.R., A.O.O., E.B., A.B.T., and J.B. contributed to experimental planning, device methodology, and operation. J.C.T., T.J.C., A.L., J.M.W., X.S., A. Shin, E.B., A.B.T., and J.B. conducted the investigation and curated the flow data. T.J.C., J.C.T., A. Shaposhnikov, J.M.W., X.S., I.H.R., and J.B. analyzed the flow and layering data. J.C.T., T.J.C., A. Shaposhnikov, A.B.T., and J.B. validated the data. J.C.T., T.J.C., A.B.T., J.B. designed and created the manuscript figures. J.C.T., T.J.C., A. Shaposhnikov, C.S.C.L., A. Shin designed and milled the devices and obtained materials. J.C.T., T.J.C., T.M.N., A.O.O., A.B.T., J.B. managed and administered the study and T.M.N., A.O.O., E.B., A.B.T., J.B. advised on study design and execution. J.C.T., T.J.C., A.L., A.B.T., J.B. wrote sections of the manuscript. All authors revised and approved the manuscript.

