## Supporting Information for "Modeling and validation of parallel co-flow layer widths of viscous liquids in open-capillary trigger valve systems"

**Table of Contents**


| **Section** | | **Page** |
|---|---|---|
| SI.1 | Engineering drawings of trigger valve devices | S2 |
| SI.2 | Experimental travel distance data for all trials used in this study | S3 |
| SI.3 | Experimental layer width data for all trials used in this study | S9 |
| SI.4 | Theory notation | S12 |
| SI.5 | Images of hydrogel patterning replicates | S14 |

### SI.1. Engineering drawings of trigger valve devices

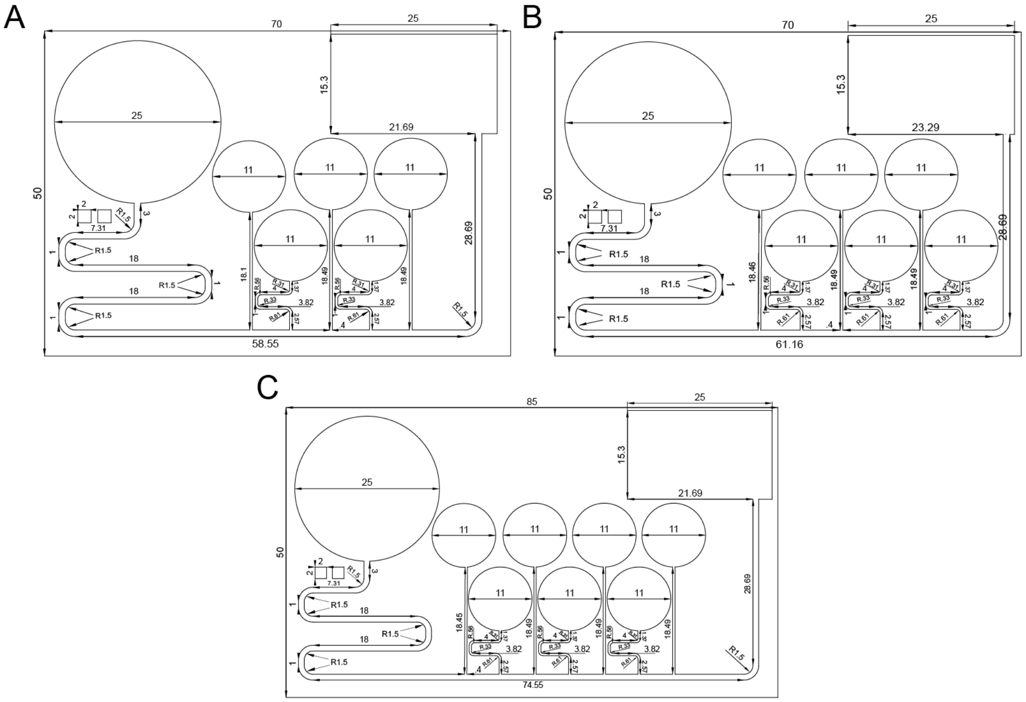


Figure SI.1.1. Engineering drawings of 5-valve (A), 6-valve (B), and 7-valve (C) trigger valve systems. Note: For the 0.6 mm and 0.7 mm deep 5-valve TGV systems, the same top-down sketch was used for both devices and only the z height of the side channels were varied. All dimensions in millimeters.

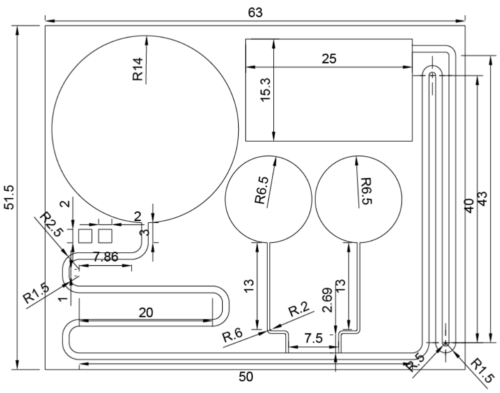


Figure SI.1.2. Engineering drawing of a 2-valve trigger valve system. Channel depth is 0.7 mm for the TGVs and 1 mm for the main channel.

**SI.2. Experimental travel distance data for all trials used in this study**

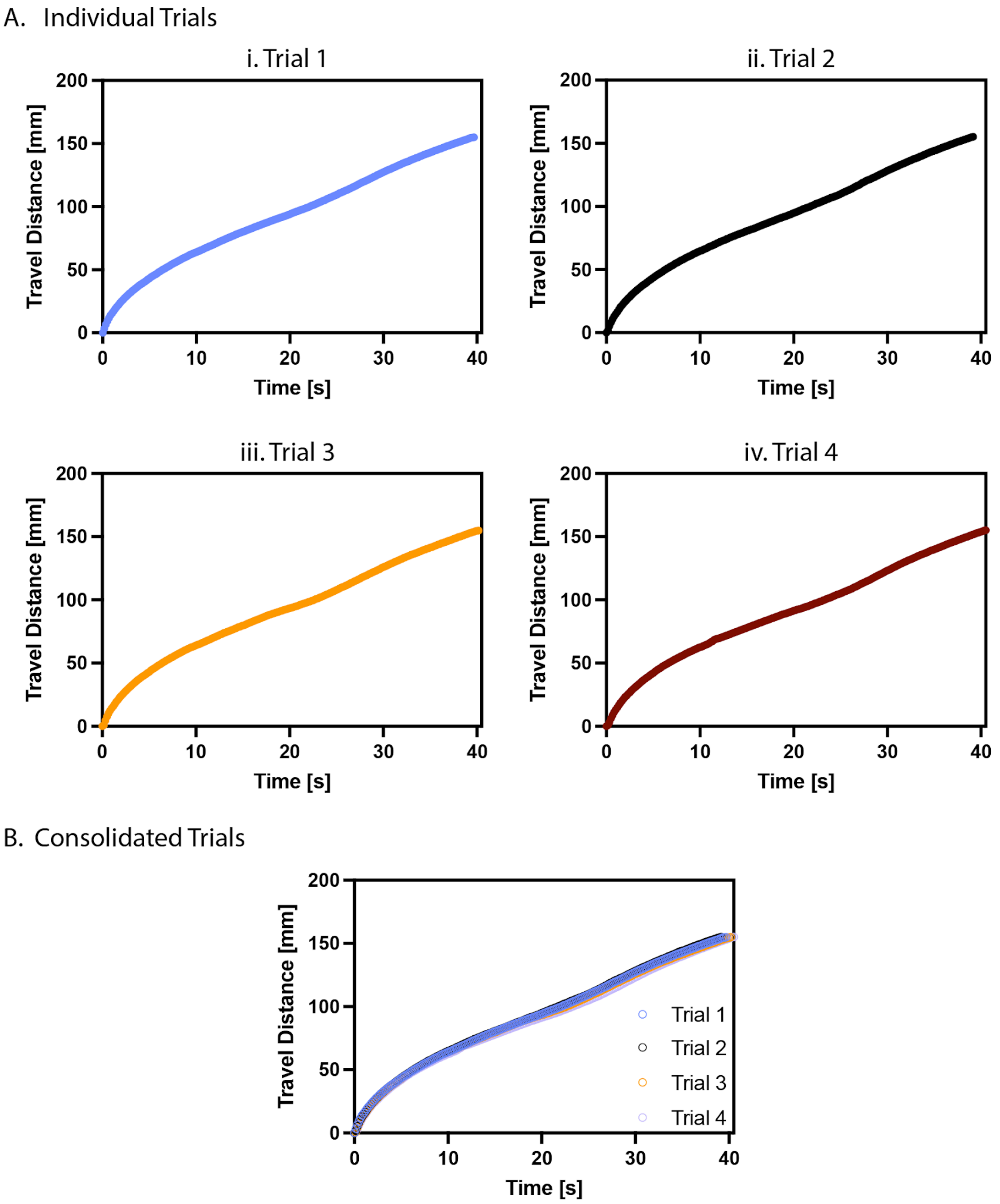


**Figure SI.2.1.** Experimental data for trials 1 to 4 (Ai to Aiv) for a 5-valve TGV system with a side channel depth of 0.6 mm. (B) All trials overlayed in a single graph.

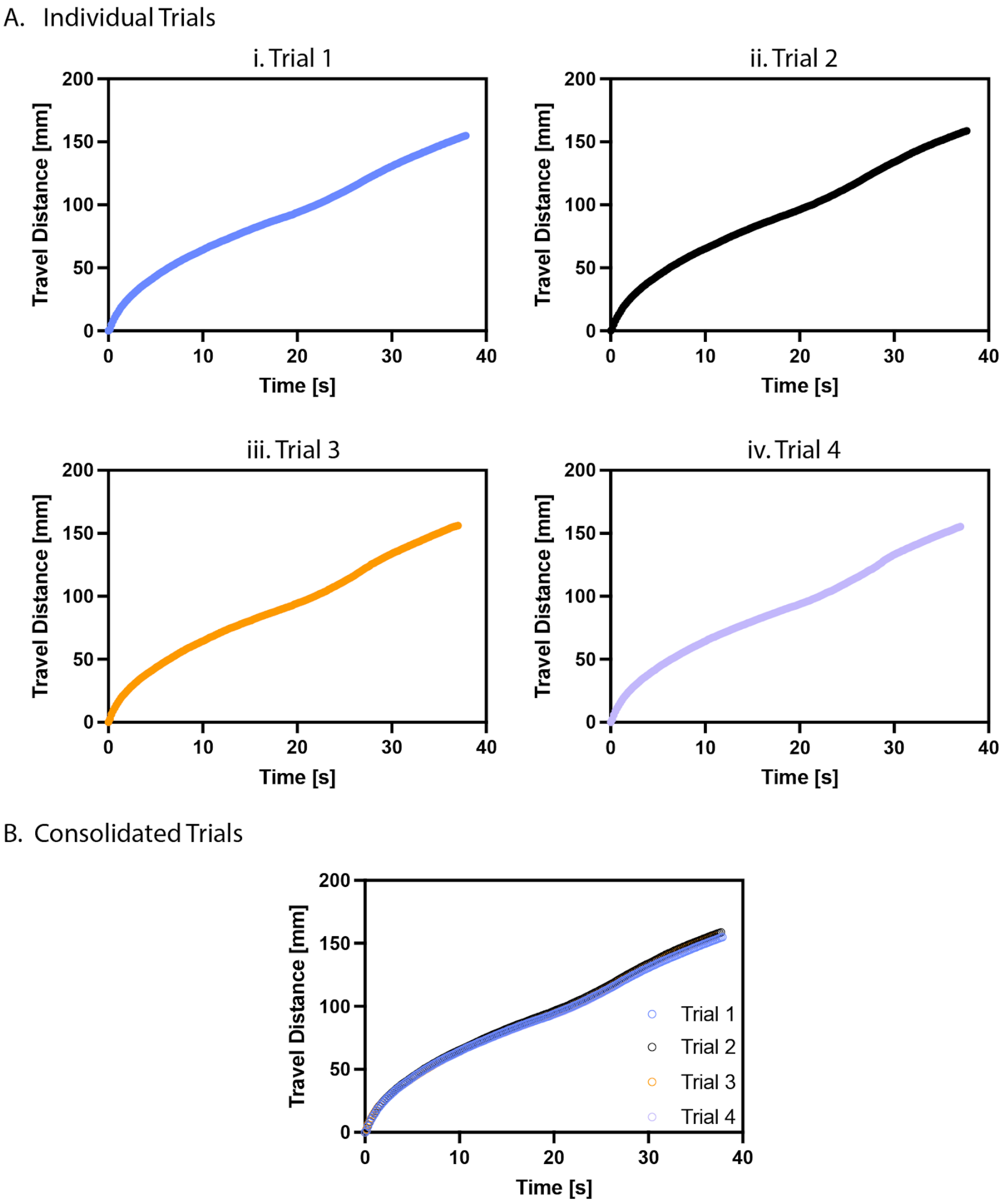


**Figure SI.2.2.** Experimental data for trials 1 to 4 (Ai to Aiv) for a 5-valve TGV system with a side channel depth of 0.7 mm. (B) All trials overlayed in a single graph.

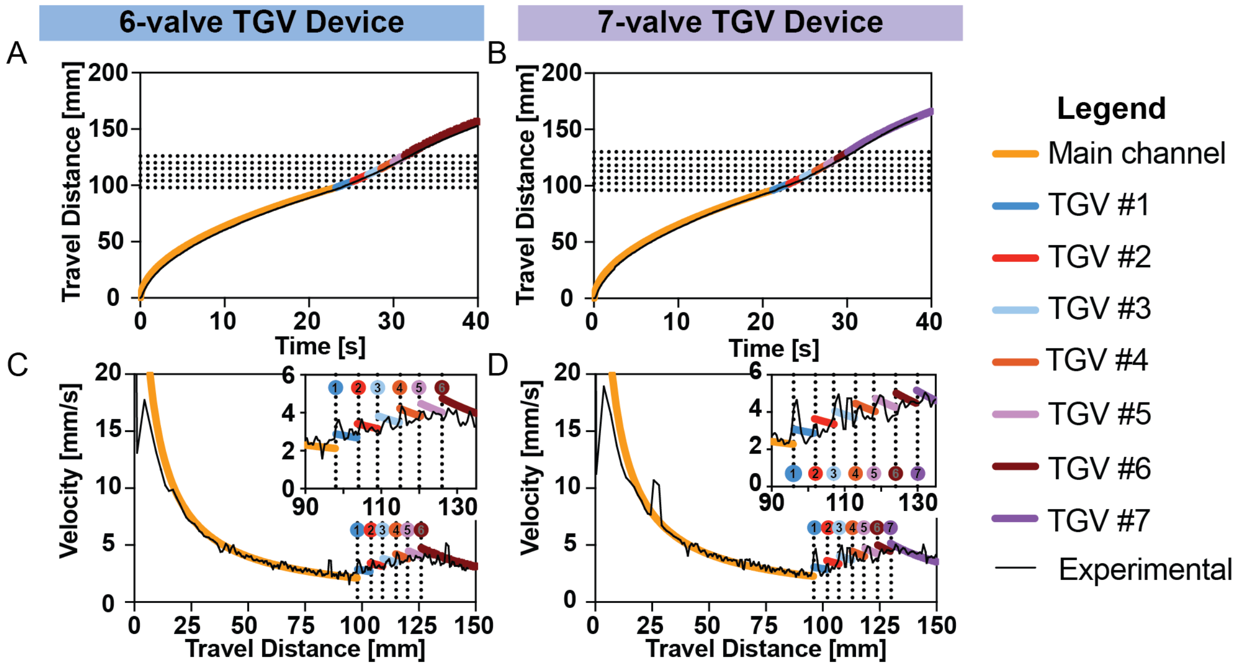


**Figure SI.2.3** Flow in devices with six and seven TGVs (A) Travel distance over time in devices with 6 TGVs. (B) Travel distance over time in devices with 7 TGVs for a representative trial. (C) Velocity over travel distance in devices with 6 TGVs for a representative trial. (D) Velocity over travel distance in devices with 7 TGVs for a representative trial. Travel distance data for n = 3 for the 6 TGVs and 7 TGVs devices shown in Figure SI.2.4 and SI.2.5, respectively.

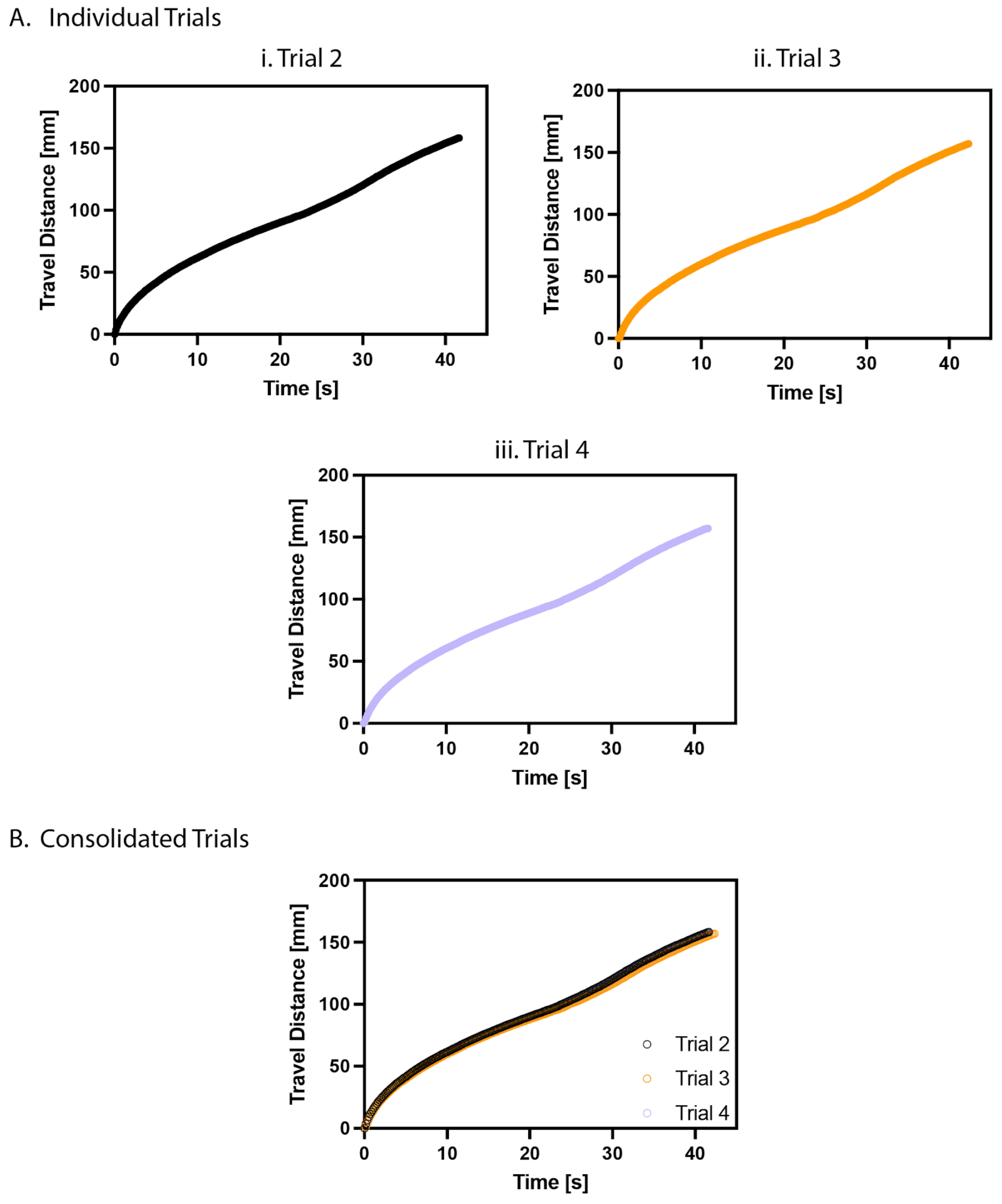


**Figure SI.2.4.** Experimental data for trials 2 to 4 (Ai to Aiii) for a 6-valve TGV system. (B) All trials overlayed in a single graph.

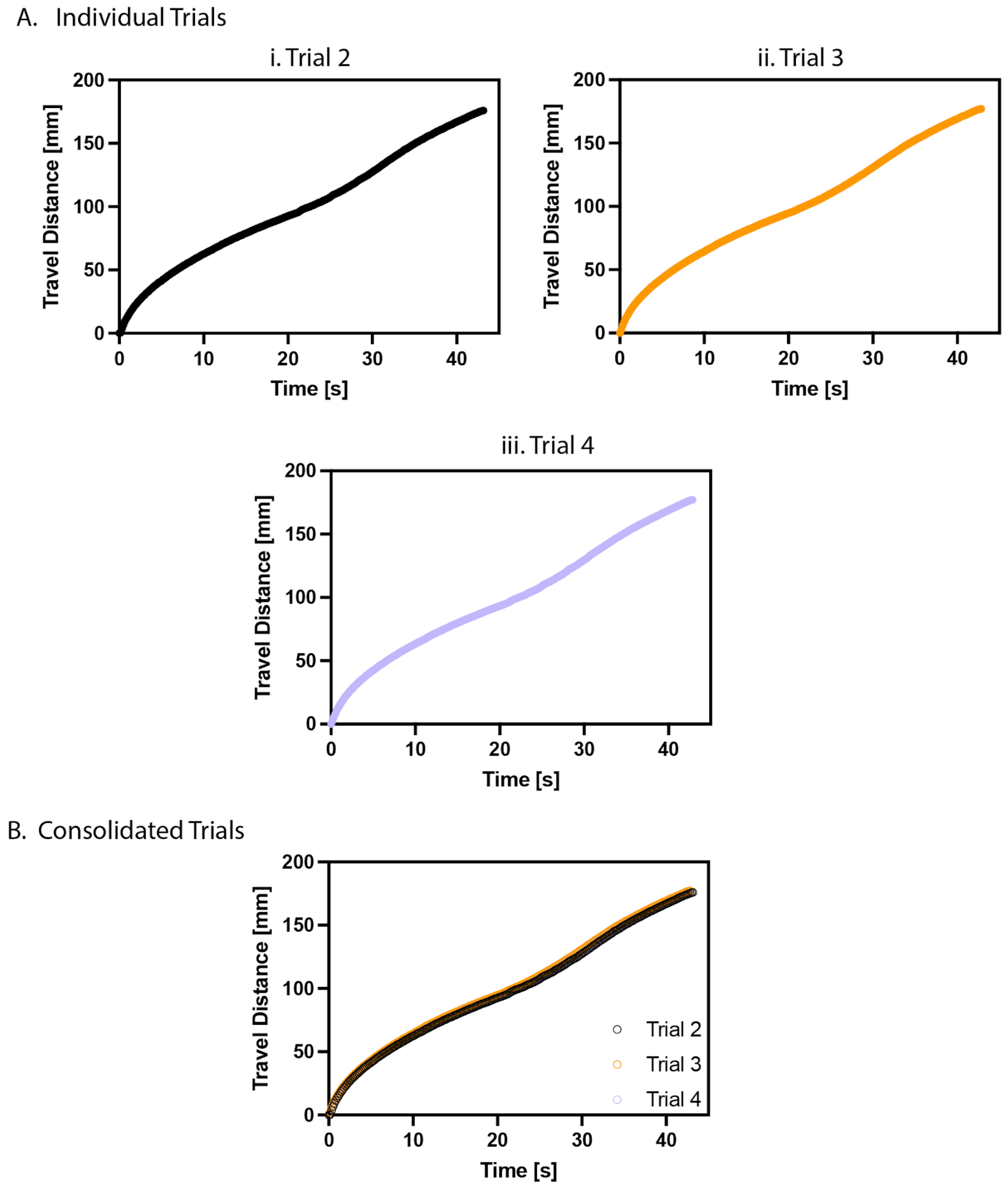


**Figure SI.2.5.** Experimental data for trials 2 to 4 (Ai to Aiii) for a 7-valve TGV system. (B) All trials overlayed in a single graph.

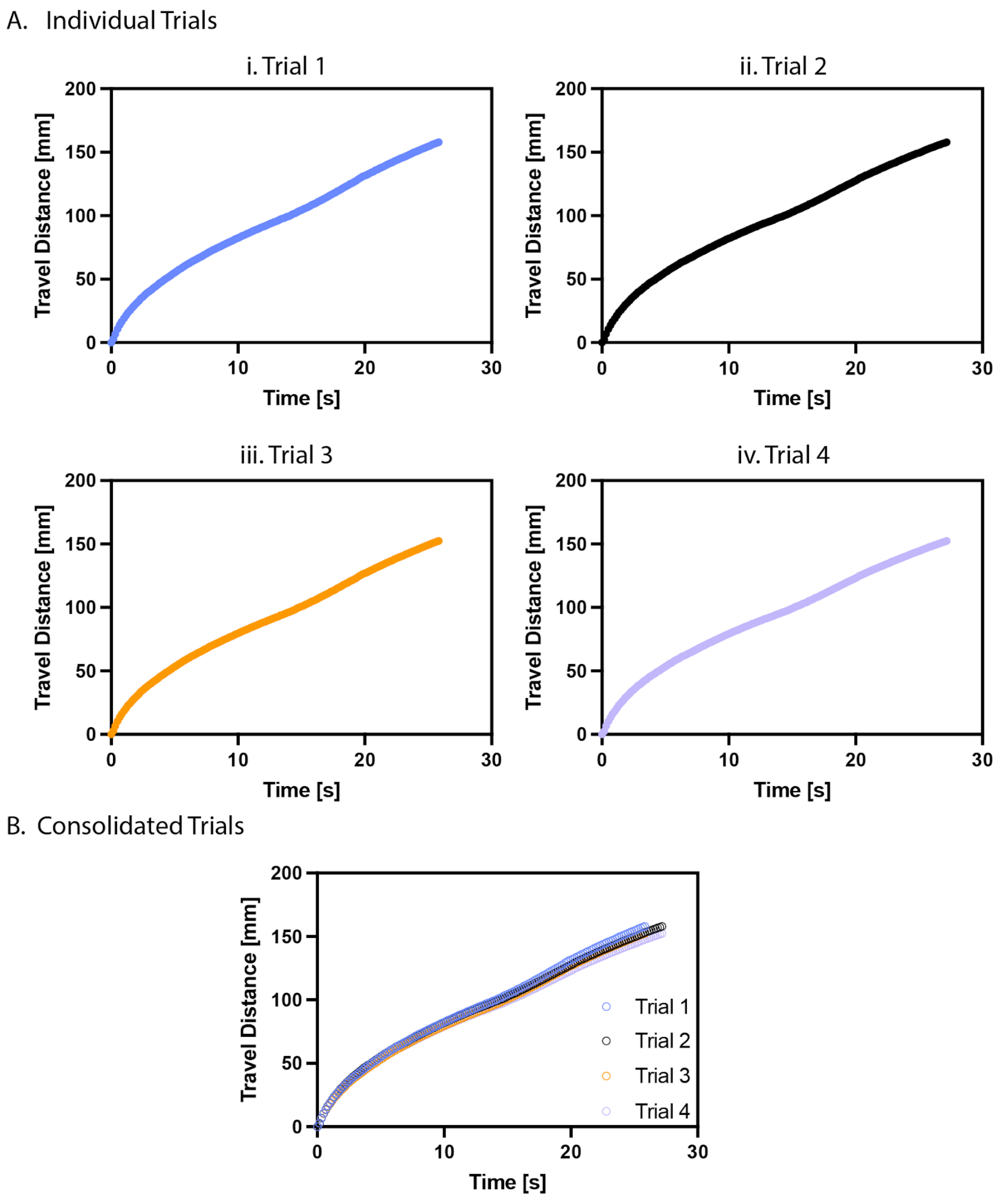


**Figure SI.2.6.** Experimental data for trials 1 to 4 for a 5-valve TGV system using a glycerol/water solution.

**SI.3. Experimental layer width data for all trials used in this study**

**Table SI.3.1**. Layer widths determined after the release of the final valve in a 5 TGV PMMA device with 0.6 mm depths. Layer widths after each valve release is provided in the raw data file.

| | Trial 1 | Trial 2 | Trial 3 | Trial 4 | Average | Standard Deviation | Theory value |
|---|---|---|---|---|---|---|---|
| Main Flow (mm) | 0.335 | 0.360 | 0.349 | 0.380 | 0.356 | 0.0190 | 0.3587 |
| Layer 1 (mm) | 0.049 | 0.065 | 0.079 | 0.065 | 0.0645 | 0.0123 | 0.0890 |
| Layer 2 (mm) | 0.078 | 0.086 | 0.079 | 0.087 | 0.0825 | 0.00465 | 0.1001 |
| Layer 3 (mm) | 0.100 | 0.116 | 0.094 | 0.098 | 0.102 | 0.00966 | 0.0994 |
| Layer 4 (mm) | 0.150 | 0.120 | 0.121 | 0.124 | 0.12875 | 0.0143 | 0.1344 |
| Layer 5 (mm) | 0.275 | 0.254 | 0.255 | 0.245 | 0.25725 | 0.0127 | 0.2183 |

**Table SI.3.2.** Layer widths determined after the release of the final valve in a 5 TGV PMMA device with 0.7 mm depths. Layer widths after each valve release is provided in the raw data file.

| | Trial 1 | Trial 2 | Trial 3 | Trial 4 | Average | Standard Deviation | Theory value |
|---|---|---|---|---|---|---|---|
| Main Flow (mm) | 0.299 | 0.350 | 0.357 | 0.314 | 0.330 | 0.0280 | 0.3517 |
| Layer 1 (mm) | 0.087 | 0.063 | 0.069 | 0.082 | 0.07525 | 0.0111 | 0.0762 |
| Layer 2 (mm) | 0.101 | 0.090 | 0.092 | 0.096 | 0.09475 | 0.00486 | 0.0891 |
| Layer 3 (mm) | 0.118 | 0.102 | 0.106 | 0.126 | 0.113 | 0.0110 | 0.093 |
| Layer 4 (mm) | 0.131 | 0.143 | 0.121 | 0.136 | 0.13275 | 0.00925 | 0.1305 |
| Layer 5 (mm) | 0.261 | 0.246 | 0.237 | 0.248 | 0.248 | 0.00990 | 0.2594 |

**Table SI.3.3.** Layer widths determined after the release of the final valve in a 6 TGV PMMA device with 0.7 mm depths. Layer widths after each valve release is provided in the raw data file.

| | Trial 1 | Trial 2 | Trial 3 | Trial 4 | Average | Standard Deviation | Theory value |
|---|---|---|---|---|---|---|---|
| Main Flow (w) | 0.306 | 0.283 | 0.364 | 0.356 | 0.32725 | 0.0391 | 0.3136 |
| Layer 1 (mm) | 0.051 | 0.068 | 0.059 | 0.057 | 0.05875 | 0.00704 | 0.0675 |
| Layer 2 (mm) | 0.066 | 0.093 | 0.066 | 0.067 | 0.073 | 0.0133 | 0.0774 |
| Layer 3 (mm) | 0.080 | 0.089 | 0.081 | 0.075 | 0.08125 | 0.00580 | 0.067 |
| Layer 4 (mm) | 0.090 | 0.083 | 0.069 | 0.082 | 0.081 | 0.00876 | 0.0972 |
| Layer 5 (mm) | 0.138 | 0.143 | 0.113 | 0.117 | 0.12775 | 0.0150 | 0.1117 |
| Layer 6 (mm) | 0.247 | 0.249 | 0.229 | 0.246 | 0.24275 | 0.00925 | 0.2656 |

**Table SI.3.4.** Layer widths determined after the release of the final valve in a 7 TGV PMMA device with 0.7 mm depths.

| | Trial 1 | Trial 2 | Trial 3 | Average | Standard Deviation | Theory value |
|---|---|---|---|---|---|---|
| Main Flow (mm) | 0.283 | 0.298 | 0.296 | 0.29233333 | 0.00814 | 0.2808 |
| Layer 1 (mm) | 0.050 | 0.048 | 0.049 | 0.049 | 0.001 | 0.0592 |
| Layer 2 (mm) | 0.067 | 0.064 | 0.062 | 0.06433333 | 0.00252 | 0.0662 |
| Layer 3 (mm) | 0.070 | 0.051 | 0.059 | 0.06 | 0.00954 | 0.0656 |
| Layer 4 (mm) | 0.063 | 0.083 | 0.072 | 0.07266667 | 0.0100 | 0.0797 |
| Layer 5 (mm) | 0.098 | 0.097 | 0.113 | 0.10266667 | 0.00896 | 0.0839 |
| Layer 6 (mm) | 0.118 | 0.132 | 0.124 | 0.12466667 | 0.00702 | 0.1205 |
| Layer 7 (mm) | 0.245 | 0.230 | 0.227 | 0.234 | 0.00964 | 0.2443 |

**Table SI.3.5.** Layer widths determined after the release of the final valve in a 5 TGV PS device with 0.7 mm depths.

| | Trial 1 | Trial 2 | Trial 3 | Average | Standard Deviation | Theory value |
|---|---|---|---|---|---|---|
| Main Flow (w) | 0.373 | 0.383 | 0.397 | 0.38433333 | 0.0121 | 0.3876 |
| Layer 1 (mm) | 0.085 | 0.073 | 0.075 | 0.07766667 | 0.00643 | 0.0875 |
| Layer 2 (mm) | 0.086 | 0.091 | 0.091 | 0.08933333 | 0.00289 | 0.0991 |
| Layer 3 (mm) | 0.087 | 0.092 | 0.094 | 0.091 | 0.00361 | 0.0987 |
| Layer 4 (mm) | 0.144 | 0.139 | 0.125 | 0.136 | 0.00985 | 0.1327 |
| Layer 5 (mm) | 0.211 | 0.196 | 0.186 | 0.19766667 | 0.0126 | 0.1943 |

### SI.4. Theory Notation

**Table SI.4.1.** Notation used for all equations used in theory calculations.

| Name | Notation | Unit | Remarks / references |
|---|---|---|---|
| Pressure at node n | $P_n$ | mPa | Pressure at node n |
| Capillary pressure | $P_{cap}$ | mPa | Laplace pressure of the meniscus |
| Dynamic viscosity | $\mu$ | mPa·s | Liquid viscosity, nonanol |
| Velocity, main channel | $\mathrm{V}$ | mm/s | Cross-sectional average velocity, main channel |
| Velocity, side channel n | $\mathrm{V}_n$ | mm/s | Cross-sectional average, side channel n |
| Velocity, layer n | $V_n$ | mm/s | Local velocity in layer n |
| Friction length, main channel | $\lambda$ | mm | Average friction length, main channel |
| Friction length, side channel n | $\lambda_n$ | mm | Average friction length, side channel n |
| Total perimeter | $p$ | mm | Total perimeter of a cross-section |
| Perimeter, side channel n | $p_n$ | mm | Side channel n |
| Cross-sectional area, main channel | $S$ | mm² | Main channel |
| Cross-sectional area, side channel | $S_n$ | mm² | Side channel n |
| Length, side channel n | $L_n$ | mm | Side channel length |
| Axial coordinate, meniscus | $z$ | mm | Location of the advancing meniscus |
| Node location | $z_n$ | mm | Axial location of node n |
| Flow rate, side channel n | $\mathrm{Q}_n$ | mm³/s | Average flow rate, side channel n |
| Flow rate between nodes | $\mathrm{Q}_{n-1}$ | mm³/s | Between nodes n−1 and n |
| Flow rate, layer feed | $Q_{side,n}$ | mm³/s | Flow rate of side channel into layer n |
| Layer depth | $h$ | mm | Approximately the side channel depth |
| Main channel width | $w$ | mm | Width of the main channel |
| Layer width | $w_n$ | mm | Width of fluid layer n |
| Side channel width | $w_{n,s}$ | mm | Width of side channel n |
| Layer width, iterations | $w_n^{(1)}, w_n^{(2)}$ | mm | Successive iterative approximations |
| Layer velocity, iteration | $V_n^{(1)}$ | mm/s | Iterative approximation |
| Wall-normal coordinate | $y$ | mm | Perpendicular to the wall, y = 0 at the wall |
| Layer boundary coordinates | $y_{1,n}, y_{2,n}$ | mm | Boundaries of layer n |
| Index | $n$ | — | Node, side channel, or layer index |

Important Relations:

Average friction length = $\bar{\lambda} = \frac{1}{3}\,\frac{(w+h)}{\frac{w}{h}+\frac{2h}{w}}$.

Generalized Cassie angle = $\cos\theta^* = \frac{(w+2h)\cos\theta - w}{2\,(w+h)}$

Capillary force = $F = \gamma\; p\; cos\theta^*$

#### SI.5. Images of hydrogel patterning replicates

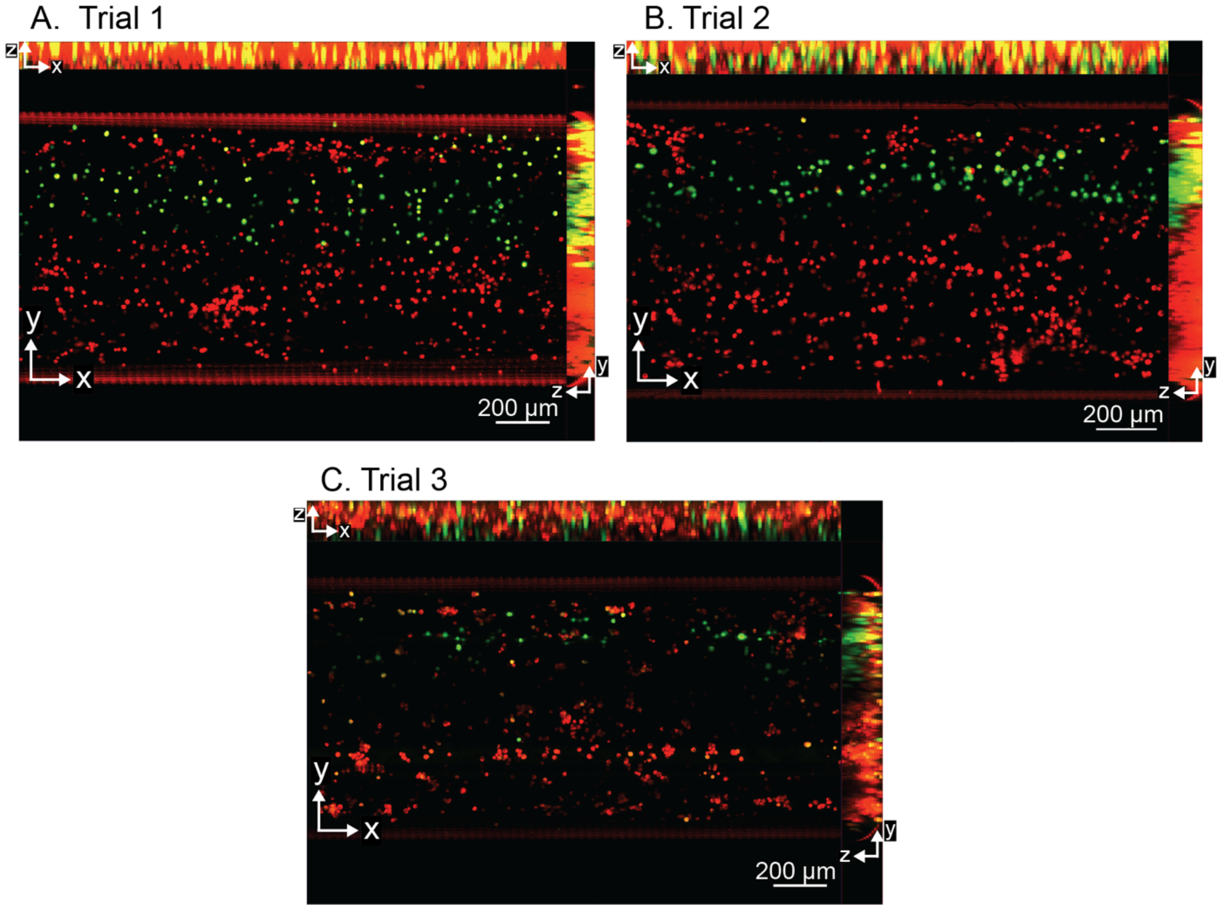


**Figure SI.5.1.** Maximum intensity projection image from bottom-up view (XY) of the main channel after the last trigger valve was released with layering of red and green fluorescent 3T3 fibroblasts of 3T3 fibroblasts in 3 mg $mL^{-1}$ collagen extracellular matrix precursor solution for trials 1 to 3 (A-C). Orthogonal projections of the layers are shown on the top (XZ) and on the right (YZ) of each image showing the z stacking of the layers. Experiment was done in three technical replicates (n=3). Part B is used in Figure 10B in the main text.
